# GIGYF2 and 4EHP Inhibit Translation Initiation of Defective Messenger RNAs to Assist Ribosome-Associated Quality Control

**DOI:** 10.1101/792994

**Authors:** Kelsey L. Hickey, Kimberley Dickson, J. Zachery Cogan, Joseph M. Replogle, Michael Schoof, Karole N. D’Orazio, Niladri K. Sinha, Adam Frost, Rachel Green, Jonathan S. Weissman, Kamena K. Kostova

## Abstract

<u>R</u>ibosome-associated <u>Q</u>uality <u>C</u>ontrol (RQC) pathways protect cells from toxicity caused by incomplete protein products resulting from translation of damaged or problematic mRNAs. Extensive work in yeast has identified highly conserved mechanisms that lead to the degradation of the faulty mRNA and partially synthesized polypeptide. Here, we used CRISPR-Cas9-based screening to search for additional RQC strategies in mammals. We found that failed translation leads to specific silencing of translation initiation on that message. This negative feedback loop is mediated by two translation inhibitors, GIGYF2 and 4EHP. Their recruitment to defective messages can be mediated by different factors, including potentially the collision sensor ZNF598. Both model substrates and growth-based assays established that inhibition of additional rounds of translation acts in concert with known RQC pathways to prevent buildup of toxic proteins. Inability to block translation of faulty mRNAs, and subsequent accumulation of partially synthesized polypeptides, could explain the neurodevelopmental and neuropsychiatric disorders observed in mice and humans with compromised GIGYF2 function.

## Introduction

Translation is an enormous amplification step in which each messenger RNA (mRNA) is turned into hundreds of proteins. Defective mRNAs lacking a stop codon due to premature polyadenylation or mRNAs harboring damage, non-optimal codons or RNA structure can cause ribosomes to stop translating before they reach a stop codon (Doma and Parker, 2006; Klauer and van Hoof, 2012; Letzring et al., 2013). Such stalled ribosomes pose numerous threats to the cells as they decrease protein output from an mRNA, block translation by trailing ribosomes, and carry incomplete polypeptide that can aggregate or exhibit dysregulated activity (Choe et al., 2016; Chu et al., 2009; Yonashiro et al., 2016). Therefore, if ribosome stalling is not detected and resolved, it can jeopardize cell and organismal viability.

To counter the threat posed by stalled ribosomes, cells have evolved multiple surveillance mechanisms, collectively referred to here as <u>R</u>ibosome-associated <u>Q</u>uality <u>C</u>ontrol (RQC) pathways. Recent studies have shown that one of the earliest events that can initiate the RQC is the collision between the stalled ribosome and a trailing one (Juszkiewicz and Hegde, 2017; Juszkiewicz et al., 2018; Simms et al., 2017; Sundaramoorthy et al., 2017). One mechanism of detection of such collided disomes is mediated by an E3 ubiquitin ligase Hel2p (ZNF598 in human) (Brandman et al., 2012; Garzia et al., 2017; Juszkiewicz et al., 2018; Letzring et al., 2013; Matsuo et al., 2017; Saito et al., 2015; Shao et al., 2015; Sundaramoorthy et al., 2017). After detection, the stalled ribosome is dissociated by the combined action of Dom34p and Hbs1p (Izawa et al., 2012). The 60S ribosomes, which still carries the stalled nascent polypeptide, recruits the core RQC components, allowing the nascent polypeptide to be ubiquitinated and extracted from the exit tunnel for proteasomal degradation (Bengtson and Joazeiro, 2010; Brandman et al., 2012; Kostova et al., 2017; Osuna et al., 2017).

These quality control pathways have been discovered and extensively studied in yeast. However, all major RQC factors are conserved from yeast to human. In contrast to yeast, where loss of core RQC factors is well tolerated under standard growth conditions, mutations in the mammalian factors affect cell growth and have been associated with complex disease. For example, partial loss of function mutations in LTN1, the ubiquitin ligase facilitating the degradation of the stalled nascent polypeptide, cause neurodegeneration in mice (Bengtson and Joazeiro, 2010; Chu et al., 2009). Together, these observations further emphasize the importance of RQC for maintaining protein homeostasis in higher eukaryotes (Brandman et al., 2012; Defenouillère et al., 2013; Shao and Hegde, 2014; Shao et al., 2013; Verma et al., 2013). Given the toxicity of partially synthesized proteins, it is possible that multiple layers of regulation exist to prevent, detect, and cope with ribosome stalling. However, our understanding of how the RQC cooperates with other pathways governing protein homeostasis is incomplete. Here we use reporter-based and growth-based genome-wide CRISPRi screens (Gilbert et al., 2014; Qi et al., 2013) to systematically characterize the mammalian RQC pathway. We characterize a new branch of the RQC that involves two factors, GIGYF2 and 4EHP, that block ribosome initiations on problematic mRNAs.

## Results

### CRISPRi screen for mammalian factors that stabilize Non-Stop decay substrate

To systematically explore how mammalian cells cope with problematic mRNAs, we designed a mammalian RQC reporter that contains an open-reading frame encoding for the green fluorescent protein, but no stop codon (GFP_Non-stop_). Incomplete proteins generated from such messages are model substrates for RQC-mediated degradation in yeast (Bengtson and Joazeiro, 2010) but have not been studied in mammals. Since the encoded GFP lacks a stop codon, translation through the GFP to the end of the message (Fig. S1A) causes ribosome stalling and subsequent degradation of the GFP nascent polypeptide via the RQC pathway. Upstream of the GFP, the reporter encodes BFP separated by a T2A ribosome skipping sequence (Fig. 1A). As a result, BFP synthesis is uncoupled from RQC-mediated degradation of GFP, allowing BFP to serve as control for the expression levels of the reporter. Since messages lacking a stop codon are rapidly degraded by the exosome (Frischmeyer et al., 2002) we introduced a triple helix derived from the MALAT1 non-coding RNA at the 3’ end to stabilize the message (Wilusz et al., 2012). Wild type cells accumulate BFP, but not GFP, resulting in a substantial decrease (~ 100 fold) in the GFP/BFP ratio for the GFP_Non-stop_ reporter compared to two control reporters that contain either a stop codon before the MALAT1 sequence (GFP_stop_), or a stop codon and a canonical polyA tail (GFP_polyA_) (Fig. 1A, B). Knockdown of a core RQC component, *NEMF,* led to the stabilization of GFP (Fig. 1C, Fig. S1B), confirming the role of mammalian NEMF in degradation of nascent polypeptides resulting from non-stop decay mRNAs (Shao et al., 2015).

**Fig. 1.**
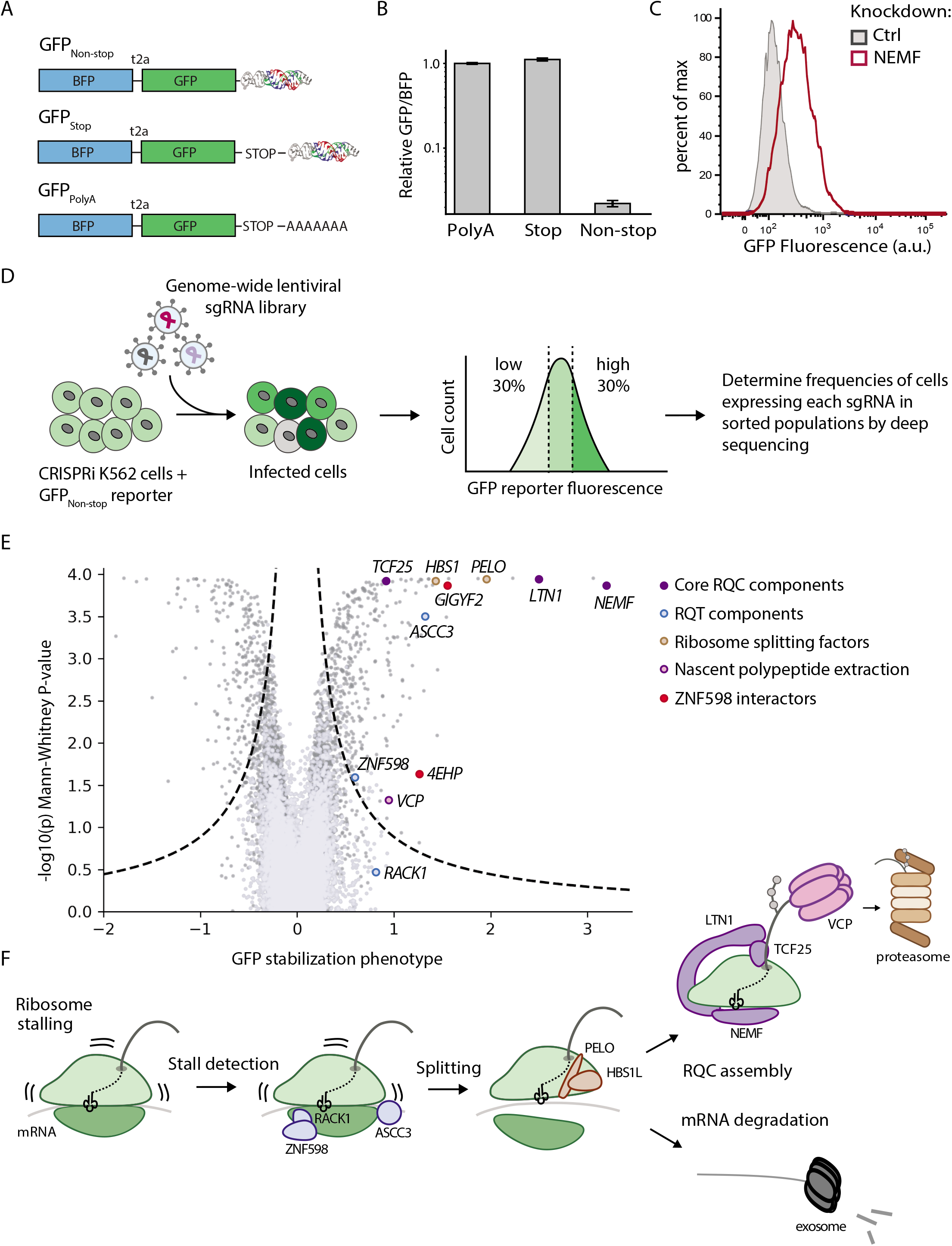
Genome-wide CRISPRi screen for mammalian RQC components. (**A**) Diagrams of non-stop stalling reporter (GFP_Non-stop_) and control reporters (GFP_Stop_ and GFP_polyA_) (**B**) Median GFP:BFP ratio of 293T cells transiently transfected with reporter constructs containing the indicated 3’ mRNA sequence (median ± SD, N=3). (**C**) GFP fluorescence level in cell lines stably expressing the GFP_Non-stop_ reporter and control sgRNA or sgRNA against *NEMF.* (**D**) Workflow of FACS-based CRISPRi screen. Reporter cell line constitutively expressing GFP_Non-stop_ is infected with the whole genome CRISPRi library. Knockdown of genes involved in coping with ribosome stalling leads to GFP accumulation. GFP positive cells are sorted out and the sgRNAs expressed in those cells are identified via deep sequencing. (**E**) Volcano plot of GFP stabilization phenotype and Mann-Whitney P-values from genome-scale CRISPRi FACS screen. Negative controls are shown in lavender, targeting guides in grey, and previously characterized RQC factors are labeled with unique colors. (**F**) Model of the mammalian RQC pathway with screen hits highlighted.

We next used the GFP_Non-stop_ reporter in a FACS-based genome-wide CRISPRi screen to gain a comprehensive view of mechanisms that cells have for minimizing the accumulation of nascent polypeptides resulting from mRNAs lacking a stop codon. We engineered a mammalian cell line that constitutively expresses the GFP_Non-stop_ reporter and dCas9-KRAB inhibitory system (Gilbert et al., 2014). We infected this cell line with sgRNA library (V2 CRISPRi) targeting each known protein coding open reading frame (Horlbeck et al., 2016). We hypothesized that depletion of RQC factors will interfere with the ability of the cells to detect stalled ribosomes and degrade the stalled nascent polypeptide, which will result in GFP stabilization. We then sorted the cells with high GFP signal via fluorescent activated cell sorting (FACS) and identified the genes that were depleted in those cells by deep sequencing the sgRNAs they expressed (Fig. 1D, E). As expected, the vast majority of known components from each stage of the RQC pathway (stall detection, ribosome splitting, nascent chain extraction, mRNA and peptide degradation) were among the top hits (Fig. 1E, F, S1C, D).

### Modifier screen identifies the translation inhibitors GIGYF2 and 4EHP as RQC components working in parallel to nascent polypeptide degradation by NEMF

In addition to factors previously implicated in the RQC pathway, the FACS-based screen yielded several genes with no known RQC-related function. To differentiate between factors that allow mammalian cells to cope with stalled ribosomes as part of the RQC from components of parallel proteostasis pathways, we complemented our reporter-based screen with a growth-based CRISPRi modifier screen. We aimed to search for components that have synergistic growth defects in combination with loss of NEMF. This idea was motivated by recent findings that the NEMF homolog in bacteria (RqcH) leads to a synergistic growth defect when combined with loss of the tmRNA/ ssrA pathway, which helps dispose of incomplete translation products (Lytvynenko et al., 2019). To test whether similar pathways exist in combination with the mammalian RQC pathway, we took advantage of the growth defect caused by NEMF depletion (Fig. 2A). We engineered a constitutive *NEMF* knockdown cell line, as well as a control cell line, and infected them with the genome-wide CRISPRi library (Fig. 2B). We determined the change in the sgRNA abundance between the two cell lines following 10 cell doublings. We expected that depletion of RQC components will have similar effect on growth alone or in combination with *NEMF* knockdown (buffering interaction) (Fig. 2C). On the other hand, disruption of pathways that work in parallel with RQC to prevent the accumulation of failed translation products is expected to exacerbate the growth phenotype caused by loss of *NEMF* (synergistic interaction). Indeed, the screen revealed several factors that exhibit buffering or synergistic growth interactions with loss of NEMF (Fig. 2D).

**Fig. 2.**
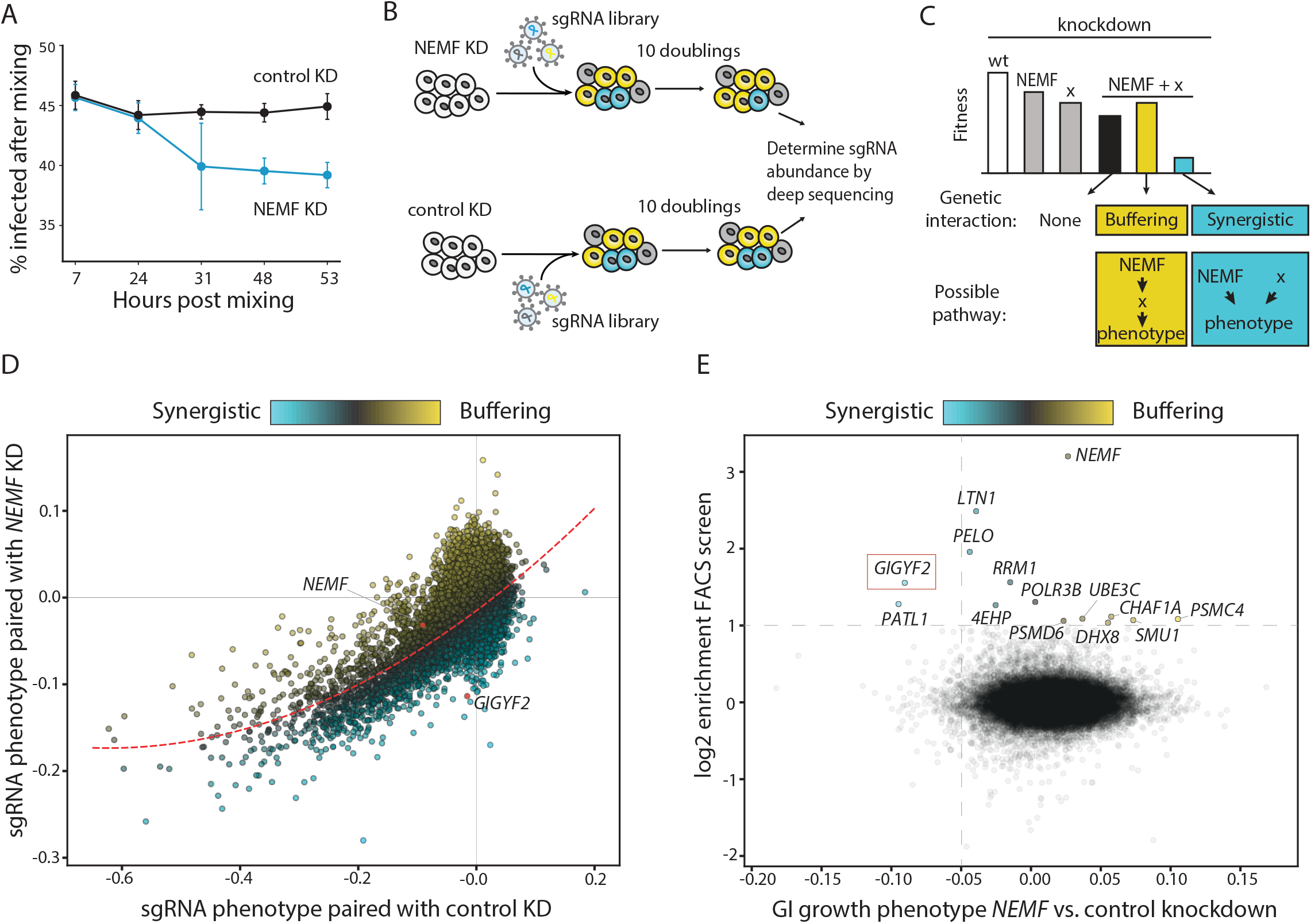
CRISPRi genetic interaction screen for factors that affect the growth of NEMF knockdown cells. (**A**) Competition assay between cells expressing a control or NEMF targeting sgRNA. (B) Workflow of growth-based CRISPRi screen. Control knockdown or NEMF knockdown cell lines were infected with the genome-scale sgRNA library and the change in the sgRNA abundance between the two cell lines following 10 cell doublings was determined by deep sequencing. (**C**) Expected growth phenotypes and predicted biological pathways resulting from knockdown of a hypothetical factor (X) alone or in combination with NEMF. (D) Results from genetic interaction (GI) screen for factors affecting the growth of control or *NEMF* knockdown cells. GI scores were derived by fitting the data to a quadratic curve (red dashed line) and calculating the distance of each point from the best-fit line. Genes exhibiting synergistic interactions with *NEMF* knockdown vs. control knockdown (negative GI score) are marked in blue, and genes with positive GI score (buffering interactions) are in yellow. (**E**) Cross comparison of FACS and GI CRISPRi screens. Hits from FACS screen (log2 enrichment > 1) that stabilized GFP_non-stop_ reporter upon individual re-testing are labeled and colored by GI score.

We then cross-compared the hits from the FACS-based reporter screen and the modifier screen. Two genes stood out as having both a strong synergistic interaction with loss of NEMF and stabilizing effect on the RQC reporter upon knockdown (Fig. 2E); PATL1 and GIGYF2. PATL1 is a characterized scaffold protein that bridges mRNA decapping and deadenylation (Ozgur et al., 2010). Stabilization of damaged mRNAs could explain both the increased levels of stalled nascent polypeptides and the synergistic growth interaction with *NEMF* knockdown, which disposes of incomplete protein products of such damaged mRNAs. We focused our attention on the second factor, GIGYF2, which has not been previously implicated in the cellular response to damaged messages.

Overexpression studies of tagged GIGYF2 have shown that it interacts with the inhibitory cap-binding protein 4EHP (Peter et al., 2017) and the ribosome collision sensor ZNF598 (Morita et al., 2012; Tollenaere et al., 2019). We confirmed the presence of a stable interaction among GIGYF2, 4EHP and ZNF598, by endogenously tagging GIGYF2 with GFP11 (Leonetti et al.) and performing co-immunoprecipitation experiments (Fig. 3A). In addition, mass spectrometry of factors that co-immunoprecipitate with FLAG-tagged ZNF598 also identified GIGYF2 and 4EHP as top interactors (Fig. S2).

**Fig. 3.**
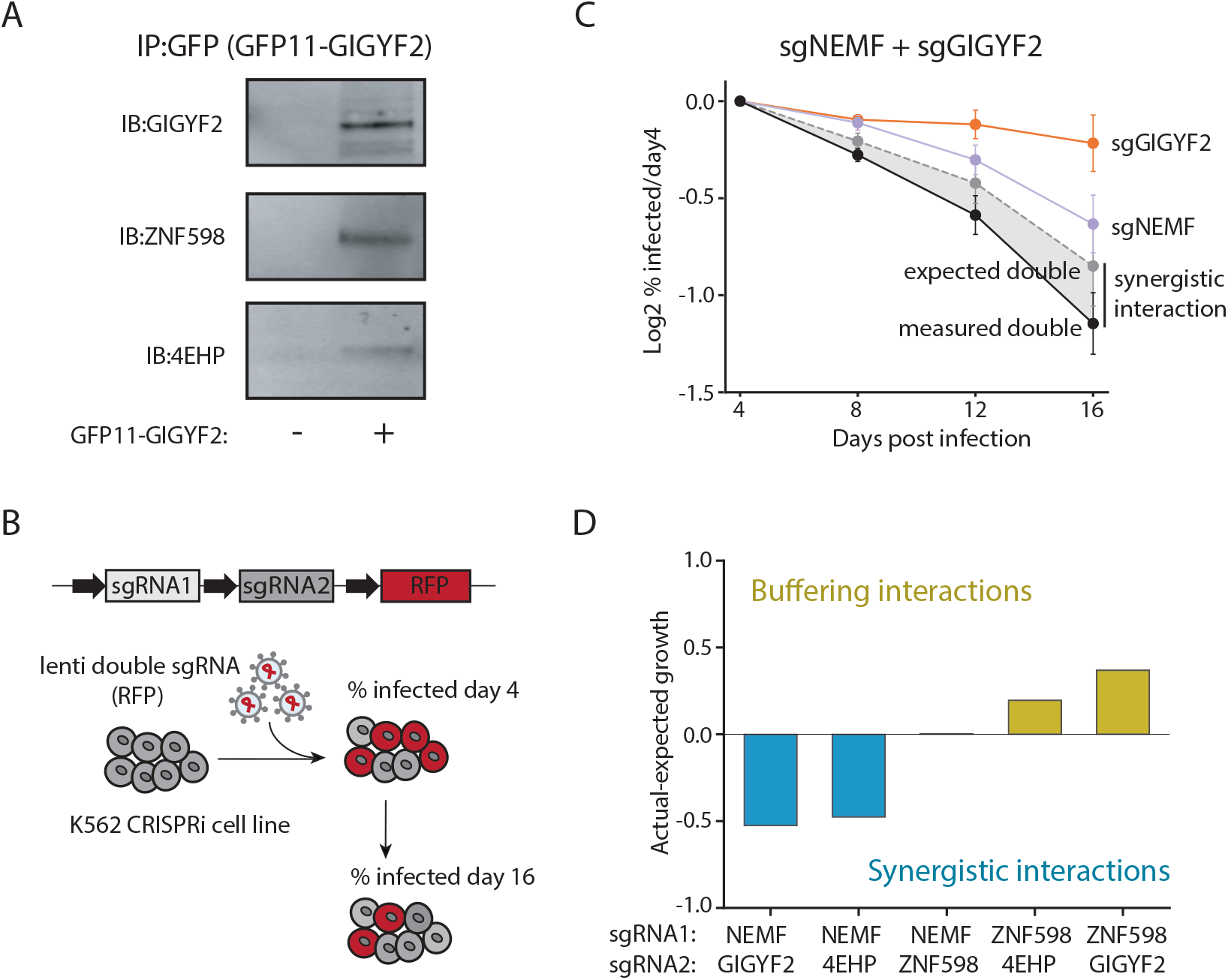
GIGFY2 and 4EHP are components of a pathway parallel to the RQC. (**A**) Coimmunoprecipitation (CO-IP) and immunoblot (IB) for endogenously tagged GIGYF2, and its binding partners, ZNF598 and 4EHP. (**B**) Outline of growth competition experiments. K562 cells were infected with RFP labeled construct carrying two sgRNAs targeting one or multiple genes. The abundance of RFP positive cells is measured over time via flow cytometry. (**C**) Competition assay among cells expressing sgRNA targeting GIGYF2 and NEMF alone or in combination (mean ± SD, N=2). Dotted grey line highlights the expected growth phenotype and the black line represents the observed growth defect of the double knockdown cells. The deviation from the expected phenotype is indicative of synergistic growth interaction. (**D**) Competition assay of double knockdown cell lines. The bar graph represents the difference in the measured (actual) growth defect and expected phenotype (additive value of single growth defects) on day 16 of the competition assay (mean ± SD, N=2).

Although an interaction between GIGYF2 and ZNF598 has been reported, the role of GIGYF2 and 4EHP in ribosome-associated quality control has not been explored. Based on our genetic interaction screen, knockdown of GIGYF2 or 4EHP in combination with knockdown of NEMF leads to a synergistic growth defect. This result suggests that GIGYF2 and 4EHP function in parallel to the RQC pathway to counter accumulation of toxic polypeptides resulting from ribosome stalling. We were able to recapitulate the genetic interaction (GI) screen results in targeted studies (Fig. 3B). The moderate growth phenotype imparted by knocking-down *NEMF* was exacerbated in combination with *GIGYF2* knockdown (Fig. 3C). Similarly, *4EHP* knockdown also had a synergistic interaction with *NEMF* knockdown, suggesting that GIGYF2 and 4EHP work in parallel to NEMF. In addition, *ZNF598* knockdown had buffering interactions with GIGYF2 (Fig. 3D, Fig S3) consistent with the data that these three proteins form a complex and work together in the same pathway.

### GIGYF2, 4EHP, and ZNF598 inhibit translation of faulty messenger RNAs

4EHP (EIF4E2) is an ortholog of the mRNA cap-binding and translation initiation factor EIF4E1. However, 4EHP cannot bind EIF4G and as a result it blocks assemble of productive EIF4F initiation complex (Rom et al., 1998; Zuberek et al.), leading to translational silencing of the bound mRNA. Indeed, recruitment of GIGYF2 or 4EHP to reporter messages has been shown to block ribosome initiation (Kryszke et al., 2016; Morita et al., 2012). It has been hypothesized that GIGYF2 requires an adapter protein to be recruited to its target mRNA, although the endogenous substrates and recruitment factors remain incompletely characterized. Our genetic and biochemical studies suggest a model where by ZNF598 could serve as one such factor which recruits GIGYF2 and 4EHP to messages harboring stalled ribosomes, which ultimately leads to translational silencing of the mRNA. To test this model, we measured the effect of knocking down these proteins on the GFP_Non-stop_ fluorescent reporter. We compared the levels of BFP, which is released before stalling and is, therefore, a readout of the reporter expression levels, and GFP, which is degraded due to ribosome stalling (Fig. 4A). As expected, knockdown of *NEMF,* a factor involved in the degradation of the stalled nascent polypeptide, stabilizes GFP, but not BFP. Knockdown of *GIGYF2* and *4EHP,* however, increased the levels of both fluorescent proteins (Fig. 4A) without affecting reporter mRNA levels (Fig. 4B), confirming the hypothesis that these two factors are translation inhibitors. To directly measure the changes in ribosome occupancy on our GFP_Non-stop_ stalling reporter, we performed ribosome profiling. Consistent with the fluorescent reporter assay, the translation efficiency (TE) of the reporter was increased in both GIGYF2 and 4EHP knockdown cell lines (Fig. S4A, B).

**Fig. 4.**
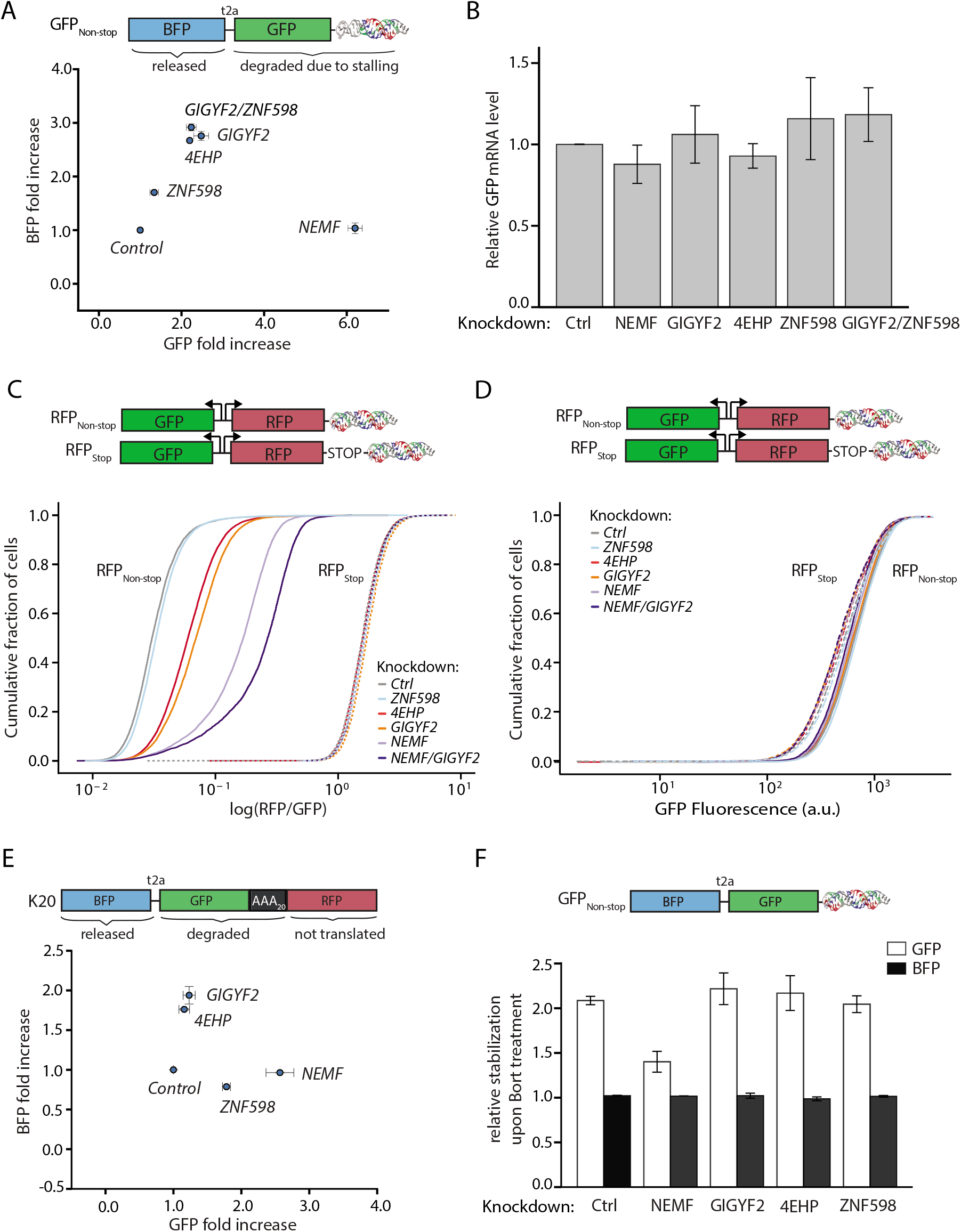
GIGYF2 and 4EHP selectively inhibit translation of faulty messages harboring stalled ribosomes. **(A)** BFP and GFP fold change upon RQC factors knockdown measured by flow cytometry (median ± SD, N=2). (**B**) Relative GFP_Non-stop_ reporter RNA levels measured by qPCR upon RQC factors knockdown (mean ± SD, N=3). (**C**) Bidirectional promoters were used to express GFP with stop codon and polyA tail, and RFP with or without stop codon (diagramed above). Cumulative distribution plot of RFP/GFP protein ratios measured by flow cytometry in RQC knockdown cells lines. RFP_Non-stop_ reporters are shown as solid lines, RFP_stop_ cell lines are shown as dashed lines. (**D**) Cumulative distribution plot of GFP protein levels measured by flow cytometry in RQC knockdown cells lines. Solid and dashed lines represent GFP signal from reporter also containing RFP_Non-stop_, and RFP_stop_ respectively. (**E**) BFP and GFP protein fold change of K20 stalling reporter upon RQC factors knockdown (median ± SD, N=2). (**F**) Cells expressing GFP_Non-stop_ reporter were treated with the proteasome inhibitor Bortezomib for 3h and the BFP and GFP protein stabilization was measured by flow cytometry (median ± SD, N=2).

To ensure that the observed translational effect was specific to faulty mRNAs, and not due to global changes in protein production, we used a second set of reporters with bidirectional promoters. One promoter drives GFP with a canonical stop codon and polyA tail, and the other drives a non-stop reporter (RFP_Non-stop_) or a control fluorescent protein (RFP_stop_) on a separate mRNA. Knockdown of *NEMF, GIGYF2,* and *4EHP* increased the RFP/GFP ratio for RFP_Non-stop_, but not RFP_stop_ (Fig. 4C). These knockdowns did not have a global effect on translation as the GFP levels were unchanged (Fig 5D). The observed RFP_Non-stop_ stabilization was not due to changes in mRNA levels (Fig S4C), consistent with the above GFP_Non-stop_ reporter. In addition, a double knockdown of *NEMF* and *GIGYF2* increased the RFP/GFP ratio to a greater extent than either single knockdown, further suggesting that these factors are components of two parallel pathways coping with ribosome stalling.

**Fig. 5.**
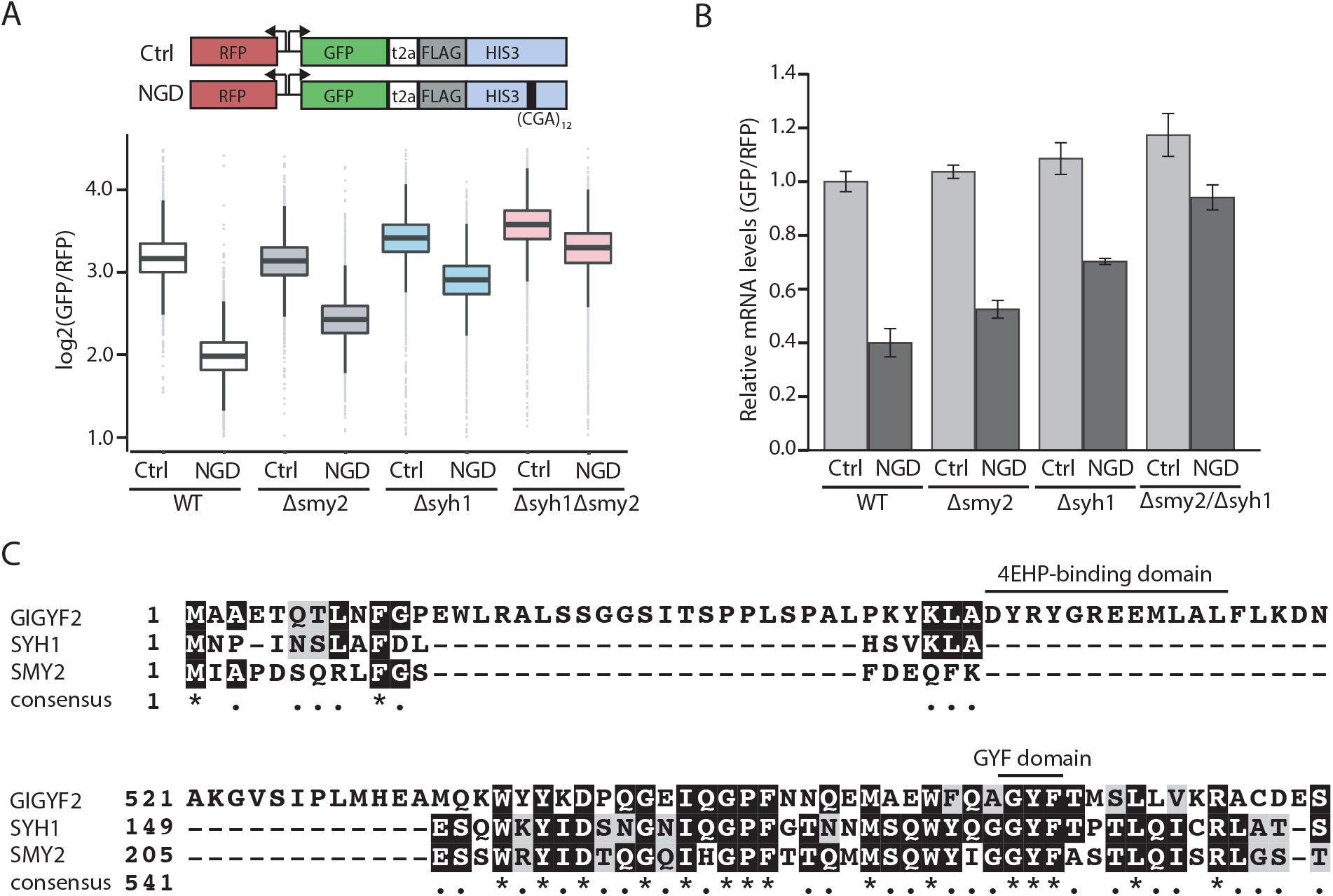
The yeast homologs of GIGYF2, Smy2p and Syh1p, are RQC factors. **(A)** Schematic of control (ctr) reporter and no-go decay (NGD) reporter containing a stretch of 12 non-optimal CGA codons in the middle of the His3 open reading frame (top). Box plot of GFP/RFP ratios for NGD or control reporter levels in knockout yeast strains measured by flow cytometry (bottom). (**B**) Relative mRNA levels for control or NGD reporter in wild type or knockout strains measured by qPCR (mean ± SD, N=3). (**C**) Alignment of regions of GIGYF2, Syh1p, and Smy2p highlighting the 4EHP-binding and GYF domains.

We confirmed that the observed translational silencing does not depend on the type of stall by measuring the effect of *GIGYF2* and *4EHP* knockdown on previously characterized No-Go reporter, containing a stretch of 20 AAA codons as a stalling sequence (K20) (Juszkiewicz and Hegde, 2017). Similar to the GFP_Non-stop_ reporter, knocking-down *GIGYF2* and *4EHP* led to an increase in both GFP and BFP fluorescence levels (Fig. 4E), without a change in mRNA levels (Fig. S4D). However, knockdown of ZNF598 led to stabilization of GFP, but not BFP. This observation is indicative of stalling-specific differences and could suggest that there are alternate factors capable of recruiting GIGYF2 and 4EHP to different classes of problematic messages. Intriguingly, prior to the discovery of the RQC pathways, it had been previously suggested that the lack of a stop codon can interfere with translation of a message (Akimitsu et al., 2007). However, the relationship (if any) between that phenomenon, which was suggested to act in a post initiation step resulting in halting of translation before completion of full length proteins (Akimitsu et al., 2007) and the GIGYF2/4EHP-mediated inhibition of translation of Non-stop and No-Go substrates is unclear.

Finally, we tested whether knockdown of GIGYF2 or 4EHP blocks RQC-mediated degradation of the stalled nascent polypeptide. We inhibited the proteasome by treating various knockdown cell lines with bortezomib (Fig. 4F). We observed further stabilization of the stalled polypeptide in the *GIGYF2, 4EHP* and *ZNF598* knockdown cells lines, but not in *NEMF* knockdown cell lines consistent with the role of *NEMF* in UPS-mediated nascent chain degradation (Fig. 3C). This result indicates that the observed increase in fluorescence upon *GIGYF2* and *4EHP* knockdown is not caused by nascent chain stabilization, but by increased ribosome engagement, and that the partially synthesized polypeptide is still targeted for degradation by RQC.

### The yeast homologs of GIGYF2, Smy2p and Syh1p, destabilize faulty messenger RNAs

We next explored whether GIGYF2 and/or 4EHP played a role in ribosome quality control in yeast. Two potential homologs of GIGYF2 have been identified in yeast, Smy2p and Syh1p (Ash et al., 2010), although they have not been implicated in RQC. To determine if Smy2p and Syh1p play a role in preventing accumulation of incomplete polypeptides, we utilized a previously characterized yeast stalling reporter (D’Orazio et al., 2019). This reporter expresses a no-go substrate and a control fluorescent protein from a bidirectional promoter. The stalling substrate encodes GFP separated by a T2A ribosome skipping sequence from HIS3 gene containing an internal stretch of non-optimal codons (CGA_12_). As a control, we used a similar reporter without the CGA12 stalling sequence. We compared the protein levels by flow cytometry and found that knockout of SMY2 and SYH1 alone led to a partial increase in the GFP/RFP ratio (Fig. 5A). Deleting both homologs led to a greater increase in GFP/RFP compared to the single deletes, suggesting that Smy2p and Syh1p have redundant function. This redundancy could account for the lack of identification of these factors in early RQC screens. In contrast to the mammalian system, the increased expression of the reporter was accompanied by mRNA stabilization (Fig. 5B). This observation is consistent with the apparent lack of a yeast homolog of 4EHP which is an essential part of the mechanism of translation inhibition in mammals. Indeed, the yeast homologs lack the 4EHP binding domain found in human GIGYF2 (Fig. 5C). However, Smy2p has been shown to bind Eap1p, a protein implicated in mRNA decapping and degradation (Ash et al., 2010). These data suggest that the role GIGYF2-like proteins play in ribosome-associated quality control preceded the last common ancestor of yeast and mammals. Recruitment of GIGYF2 leads to translational silencing in higher eukaryotes, whereas Smy2p and Syh1p mediate RNA decay in yeast.

### Recruitment of GIGYF2, 4EHP and ZNF598 leads to translation inhibition

Our genetic and biochemical data suggest that mammalian GIGYF2 and 4EHP can be recruited to problematic messages by the ubiquitin ligase ZNF598 (Juszkiewicz and Hegde, 2017; Sundaramoorthy et al., 2017). To explore this hypothesis, we used an MS2-based system to tether ZNF598, GIGYF2, or 4EHP to a reporter mRNA (Bertrand et al., 1998) (Fig. 6A). We transiently co-expressed a non-stalling reporter (GFP_MS2-stop_) harboring three MS2 stem-loops in its 5’ UTR together with MS2 binding protein (MS2BP) fusions in HEK293T cells. Consistent with published data, recruitment of GIGYF2 or 4EHP to the reporter led to inhibition of translation initiation (Kryszke et al., 2016), as evidenced by the robust decrease in GFP fluorescence without significant changes in the mRNA levels (Fig. 6B, Fig. S5A). The translation level was not strongly affected by MS2BP recruitment alone, or overexpression of a FLAG-tagged GIGYF2 protein that does not get recruited to the reporter (Fig. S5B, C), indicating that the observed translational silencing was specific for the tethering of the factors to the mRNA. Additionally, recruitment of ZNF598 to the reporter induced similar translational silencing.

**Fig. 6.**
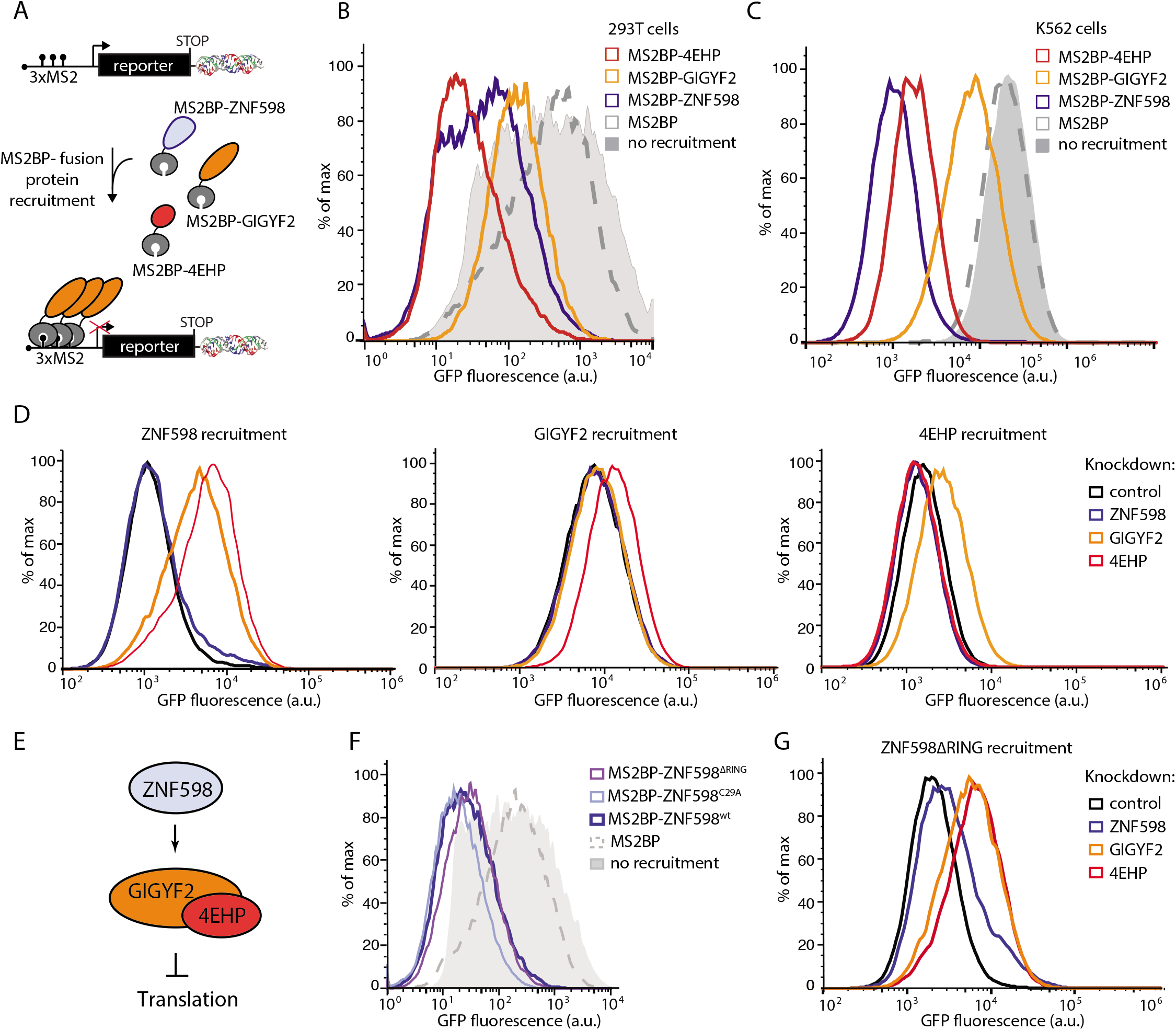
Translational inhibition by ZNF598 is mediated through GIGYF2 and 4EHP in a ubiquitination-independent manner. (**A**) Schematic of MS2-mediated recruitment of putative silencing factors to a fluorescent reporter. (**B**) GFP fluorescence of GFP_MS2-stop_ reporter transiently expressed in HEK293T cells alone or with MS2BP-fusion proteins. (**C**) GFP fluorescence of K562 cells stably expressing GFP_MS2-stop_ reporter alone or in combination with MS2BP-fusion proteins. (**D, G**) MS2-fusion protein is recruited to the GFP_MS2-stop_ reporter in control cells or knockdown cells for ZNF598, GIGYF2 or 4EHP. The effect of the knockdown on the ability of the fusion protein to silence translation from the reporter is measured via the change in GFP fluorescence. (**E**) Model for GIGYF2 and 4EHP recruitment to mRNA by ZNF598. (**F**) GFP fluorescence from GFP_MS2-stop_ reporter upon MS2-mediated recruitment of wild type or ubiquitination incompetent ZNF598 (ZNF598^C29A^ or ZNF598^ΔRING^).

To explore the genetic requirements for translation inhibition, we generated stable cell lines expressing an MS2-fusion protein, as well as GFP_MS2-stop_ reporter, and CRISPRi machinery (Fig. 6C, S5D). We used CRISPRi to knock-down *ZNF598, GIGYF2*, or *4EHP* (Fig. S7A-C) and measured the effect of the knockdown on translational inhibition via flow cytometry. We found that the decrease in GFP fluorescence caused by MS2-ZNF598 recruitment was alleviated in cells lacking GIGYF2 or 4EHP (Fig. 6D, S6D). Lower expression of the MS2BP-GIGYF2 fusion protein in K562 cells resulted in an attenuated repression phenotype compared to HEK293T (Fig. S5E). Nonetheless, the GIGYF2-mediated repression depended on 4EHP. Similarly, 4EHP-mediated repression depended on GIGYF2, suggesting that these two factors act together to mediate translation inhibition (Fig. 6D, S6E, F). In marked contrast, neither silencing by GIGYF2, nor by 4EHP were impacted by knockdown of ZNF598 when these factors were directly recruited to the mRNA, indicating that GIGYF2 and 4EHP work downstream and independently of ZNF598. These epistasis experiments support a model where ZNF598 serves as a scaffold that could recruit GIGYF2 and 4EHP to faulty mRNAs (Fig. 6E).

### Translational inhibition by ZNF598 is mediated by GIGYF2 and 4EHP in a ubiquitination-independent manner

ZNF598 recognizes collided stalled ribosomes and ubiquitinates the small subunit of the ribosome, which triggers ribosome splitting and subsequent RQC complex engagement (Ikeuchi et al., 2019; Juszkiewicz et al., 2018). We next tested whether ZNF598-mediated ubiquitination is required for translational repression by GIGYF2 and 4EHP. We generated ubiquitination-deficient MS2BP-ZNF598 fusion proteins that harbored inactivating mutations in the RING domain (C29A) or deletions of the entire domain (ΔRING) (Sundaramoorthy et al., 2017) and compared their ability to translationally silence the GFP_MS2-stop_ reporter. Surprisingly, both mutants were capable of repressing translation (Fig. 6F, S7A). Recruitment of a stably integrated MS2BP-ZNF598ΔRING also repressed translation in a GIGYF2 and 4EHP-dependent manner matching the wildtype ZNF598 recruitment. (Fig. 6G, S7B). These data suggest that ZNF598 has a dual function when it engages collided ribosomes. It serves as a ubiquitin ligase that monoubiquitinates the 40S small subunit, triggering release of the stalled ribosome and subsequent RQC engagement, and nascent polypeptide degradation (Juszkiewicz et al., 2018; Sundaramoorthy et al., 2017). In addition, ZNF598 also serves as a scaffold that provides one mechanism for the recruitment of GIGYF2 and 4EHP to the mRNA, in a ubiquitination independent manner. Once recruited, GIGYF2 and 4EHP sequester the mRNA cap, blocking ribosome initiation, and decreasing the translational load on problematic messages (Fig. 7).

**Fig. 7.**
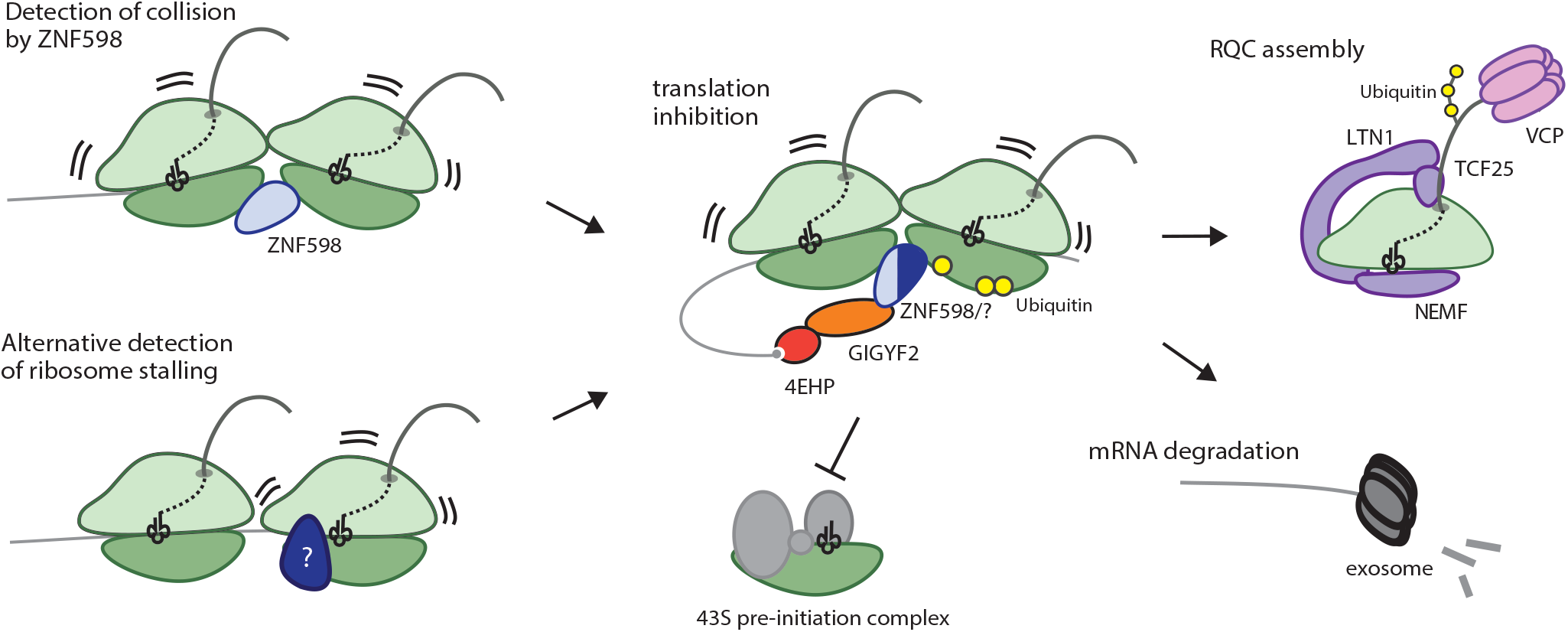
Ribosome-associated Quality Control pathway. Ribosome collision is detected by the collision sensor ZNF598. Its binding triggers a cascade of events that ultimately leads to the release of the stalled ribosome, and the degradation of the faulty mRNA and stalled nascent peptide. In addition, ZNF598 recruits the translation inhibitors GIGYF2 and 4EHP to the defective message, which blocks further ribosome initiation. Recruitment of GIGYF2 and 4EHP to defective messages could be mediated by factors other than ZNF598.

## Discussion

The cell faces an inherent challenge in detecting problematic mRNAs in that it is only through the act of translation itself that the lesion can be recognized. Our work reveals a novel mechanism in which failed translation leads to inhibition of further translation initiation on that message. This pathway acts in parallel with previously described quality control pathways, which trigger mRNA degradation, ribosome release, and proteasomal degradation of the stalled nascent polypeptide. Without a mechanism to block further initiation on problematic messages, translation of faulty mRNAs would continue for as long as the message persists. GIGYF2 and 4EHP enable the cell to break this cycle by allowing a failed translation event to initiate a specific negative feedback loop which shuts down further translation of the message. Although GIGYF2 and 4EHP have been implicated in translational control in a number of different settings (Amaya Ramirez et al., 2018; Fu et al., 2016; Kryszke et al., 2016; Morita et al., 2012; Peter et al., 2017; Tollenaere et al., 2019), our data argue that faulty mRNAs that cause ribosome stalling comprise an important subset of endogenous substrates for GIGYF2 and 4EHP mediated repression. Indeed, we see loss of GIGYF2 stabilizes two distinct classes of stalling reporters. Moreover, knocking down *GIGYF2* in combination with RQC components had a strong synergistic growth interaction in the absence of a reporter, suggesting that there is a continuous production of endogenous stalling substrates.

How then are stalled ribosomes recognized and GIGYF2 recruited? GIGYF2 and 4EHP are hypothesized to be recruited to their endogenous substrates via sets of auxiliary RNA binding proteins that serve as adapters. Our data argues that ZNF598 can serve as one of these adapters to recruit the inhibitory complex to defective messages. However, the translation phenotype of ZNF598 knockdown is consistently weaker than seen with GIGYF2 knockdown alone for Non-stop substrates (Fig. 3A) and ZNF598 unlike GIGYF2 does not have a translational effect on our No-go reporter (Fig. 4E). Moreover, when GIGYF2 is knocked-down, additional depletion of ZNF598 has no effect on the increased translation of the GFP_Non-stop_ reporter (Fig. 4A). This phenomenon is recapitulated by the artificial tethering experiments using MS2; ZNF598 relies on GIGYF2 or 4EHP for translation repression, but GIGYF2 and 4EHP do not require ZNF598 for translational repression when directly recruited to the message. Taken together, these data suggest that there are both ZNF598-dependent and ZNF598-independent mechanisms for recruitment of GIGYF2 and 4EHP to mediate translation inhibition on defective messages (Fig. 7). The identity of these additional adapters and whether they are redundant to ZNF598 or serve a specific class of faulty mRNAs remains to be explored.

Regardless of their recruitment mechanism, our data establishes a critical role for GIGYF2 and 4EHP in inhibiting ribosome initiation on defective mRNAs. The synergistic growth defects with core RQC components highlights the importance of these two factors in preventing accumulation of incomplete proteins. Loss of GIGYF2 has been associated with neurodegenerative and neurodevelopmental phenotypes (Krumm et al., 2015; Thyme et al., 2019), similar to loss of RQC components (Chu et al., 2009). It is tempting to speculate that dysregulation of the GIGYF2/4EHP pathway increases the burden on the proteostasis network in neurons. As these cells age, they can either accumulate defective mRNAs or become less efficient in detecting and/or coping with stalled ribosomes. If these cells lack functional GIGYF2 and 4EHP to translationally silence such defective messages, the increased stalling burden may overwhelm the RQC pathway in cells leading to cell stress and even death.

## Acknowledgements

We thank M. Jost, J. Hussmann, M. Shurtleff, and A. Mizrak for scientific input and helpful discussions. We thank M. Jost, and J. Hussmann for help with computational analysis. We thank R. Pak and P. Solomon for help with genetic screens. This work was supported by HHMI (J.S.W.), NIH Center for RNA Systems Biology (J.S.W.), NIH Directors’ Early Independence Award (K.K.), UCSF Genentech Fellowship (K.L.H.), HHMI Faculty Scholar award (A.F.), and NIH grant NIH 5R01AG041826-05 (J.S.W.). A.F. is a Chan Zuckerberg Biohub investigator.

## Author Contributions

K.K., K.D., K.L.H., A.F., and J.S.W. conceived the study. K.L.H., K.K. and J.S.W. wrote the manuscript with input from all authors. K.L.H., K.K., K.D., J.Z.C., M.S., K.N.D., and N.K.S. performed all experiments. J.M.R. performed ribosome profiling analysis.

## Supplementary materials

Materials and Methods

Figs. S1 to S7

Table S1-7

