## Supplemental Figures and Tables for "GIGYF2 and 4EHP Inhibit Translation Initiation of Defective Messenger RNAs to Assist Ribosome-Associated Quality Control"

**This PDF file includes:**

Materials and Methods
Figs. S1 to S7
Tables S1 to S3

**Other Supplementary Materials for this manuscript includes the following:**

Table S4: CRISPRi FACS screen gene level table
Table S5: CRISPRi growth screen gene level table

**Materials and Methods:**

Cell culture

K562 cells were grown in RPMI-1640 with 25mM HEPES, 2.0 g/L $NaHCO_3$, 0.3 g/L L-Glutamine supplemented with 10% FBS, 2 mM glutamine, 100 units/mL penicillin and 100 µg/mL streptomycin. HEK293T cells were grown in Dulbecco's modified eagle medium (DMEM) in 10% FBS, 2 mM glutamine, 100 units/mL penicillin and 100 µg/mL streptomycin.

Generation of cell lines:

To generate the K562 cell line stably expressing dCas9-KRAB, WT K562 cells were stably transduced with a lentiviral vector expressing dCas9-BFP-KRAB from an EF1-a promoter with an upstream ubiquitous chromatin opening element (SFFV-dCas9-BFP-KRAB, Addgene #85969) and selected for BFP-positive cells using two rounds of fluorescence activated cell sorting (FACS) on a BD FACSAria2.

Reporter cell lines ($GFP_{Non\text{-}stop}$, $GFP_{Stop}$, $GFP_{PolyA}$, K20, $RFP_{Non\text{-}stop}$, $RFP_{Stop}$, $GFP_{MS2\text{-}stop}$, MS2BP, MS2BP-ZNF598, MS2BP-GIGYF2, MS2BP-4EHP) were stably transduced using Piggy-Bac transposition (System Biosciences PB210PA-1 and corresponding Piggy-Bac expression vector Table 1). Positive cells were isolated by FACS on a BD FACSAria2.

Transient expression of reporters ($GFP_{Non\text{-}stop}$, $GFP_{Stop}$, $GFP_{PolyA}$, MS2BP, MS2BP-ZNF598, MS2BP-GIGYF2, MS2BP-4EHP) was achieved by transfecting HEK293T cells with vectors from Table 1 using TransIT-LTI Transfection Reagent (Mirus, MIR 2306) according to the manufacturer's instructions.

Individual gene knockdowns were carried out by selecting sgRNA protospacers from compact hCRISPRi-v2 library and cloning them into lentiviral plasmid pU6-sgRNA EF1α-puro-t2a-BFP (Addgene 60955) as previously described (Horlbeck et al., 2016). Protospacer sequences used for individual knockdowns are listed in Table 2. The resulting sgRNA expression vectors were packaged into lentivirus by transfecting HEK293T with standard packaging vectors using TransIT-LTI Transfection Reagent (Mirus, MIR 2306). The viral supernatant was harvested 2–3 days after transfection and filtered through 0.45 µm PVDF filter and/or frozen prior to transduction into CRISPRi knockdown cell lines described above.

GFP11-GIGYF2 tagged HEK293T cells were generated as previously described (Leonetti *et al.*, PNAS, 2016), using targeting sgRNA (AATACGGAACAGAATGGCAG) and ssDNA oligo(TATTTTTCTCGTTAACAGGTTTCTTCACATATAAAAATCTATTGTAAAAATACGGAAAAGAatgcgtgaccacatggtccttcatgagtatgtaaatgctgctgggattacaGGCGGTGGAGGGAGTggcggaggtGGATCCGCAGCGGAAACGCAGACACTGAACTTTGGGCCTGAATGGTGAGTTTTCAAAATCTCAT).

Genome-scale CRISPRi screening:

Genome-scale screens were conducted similar to previously described screens (Gilbert et al., 2014; Horlbeck et al., 2016). The CRISPRi compact library (5 sgRNA/TSS) CRISPRi-v2 (Addgene, Cat#83969) were transduced in duplicate into K562 CRISPRi cells at MOI < 1 (percentage of transduced cells 2 days after transduction: 20%–40%). Replicates were maintained separately in 1.5 L of RPMI-1640 in 3 L spinner flasks for the course of the screen. 2 days after transduction, the cells were selected with 1 mg/mL puromycin for 2 days, at which point transduced cells accounted for 80%–95% of the population. Cells were allowed to recover to > 80% cell viability, as measured on an Accuri bench-top flow cytometer (BD BioSciences). The cells were maintained in spinner flasks by daily dilution to $0.5 \cdot 10^6$ cells /mL at an average coverage of greater than 1000 cells per sgRNA for the duration of the screen.

*FACS based screen*

Cells were sorted using BD FACS Aria2 2 days after recovery based on GFP fluorescence of $GFP_{Non-stop}$ reporter. Cells with the highest (~30%) and lowest (~30%) GFP expression were collected and frozen after collection. Approximately 30 million cells were collected per group. Genomic DNA was isolated from frozen cells, and the sgRNA-encoded regions were enriched, amplified, and prepared for sequencing as described previously (Gilbert et al., 2014).

*Growth based Screen*

Prior to CRISPRi-v2 library infection, K562 CRISPRi cells were infected with control sgRNA or sgRNA targeting NEMF that expressed RFP as a marker. RFP positive cells were sorted on a BD FACSAria2.

Sequencing reads were aligned to the CRISPRi v2 library sequences, counted, and quantified using the Python-based ScreenProcessing pipeline (https://github.com/mhorlbeck/ScreenProcessing; Horlbeck et al., 2016). Generation of negative control genes and calculation of phenotypes and Mann-Whitney p-values was performed as described previously (Gilbert et al., 2014; Horlbeck et al., 2016). GFP phenotypes were calculated from the top 30% sorted samples, divided by the bottom 30%. Phenotypes from sgRNAs targeting the same gene were collapsed into a single GFP stabilization phenotype using the average of the top three scoring sgRNAs (by absolute value) and assigned a p-value using the Mann-Whitney test of all sgRNAs targeting the same gene compared to the non-targeting controls. All additional CRISPR screen data analyses were performed in Python 2.7 using a combination of Numpy (v1.12.1), Pandas (v0.17.1), and Scipy (v0.17.0). For all experiments, details of quantification and statistical methods used are described in the corresponding figure legends or results sections. Gene-level phenotypes are available in Tables S4-S5.

Individual evaluation of sgRNA phenotypes:

For individual evaluation and re-testing of sgRNA phenotypes, sgRNA protospacers targeting NEMF, GIGYF2, EIF4E2, or ZNF598 or a non-targeting control protospacer (neg_ctrl-1) were individually cloned by annealing complementary synthetic oligonucleotide pairs (Integrated DNA Technologies) with flanking BstXI and BlpI restriction sites and ligating the resulting double-stranded segment into either BstXI/BlpI-digested pCRISPRia-v2 (marked with a puromycin resistance cassette and RFP similar to Horlbeck et al., 2016) or BstXI/BlpI-digested pU6-sgRNA EF1a-puro-t2a-mCherry (marked with a puromycin resistance cassette and RFP). Protospacer sequences used for individual evaluation are listed in Table 2. The resulting sgRNA expression vectors were individually packaged into lentivirus. These sgRNA vectors were transduced into K562-dCas9 cells expressing $GFP_{Non\text{-}stop}$ at MOI < 1 (20 – 40% infected cells). BFP and GFP protein levels were measured by flow cytometry at day 5 post infection. Median fluorescent values were calculated for each cell line using FlowJo software and compared to uninfected levels.

Internally controlled growth assays:

sgRNA and double sgRNA growth phenotypes were performed by transducing cells with sgRNA expression constructs (mU6-sgRNA1 hU6-sgRNA2-puro-t2a-mCherry) at MOI < 1 (15 – 30% infected cells). The fraction of sgRNA-expressing cells was measured from 4-16 days after infection by flow cytometry on an LSR-II (BD Biosciences). A population of infected cells was selected to purity with puromycin (1 mg/mL), allowed to recover for 1 day, and harvested for measurement of mRNA levels by RT-qPCR (see below). Experiments were performed in duplicate from the infection step. The expected double phenotype score was calculated by adding the phenotype of both genes on day 16.
Epistasis was measured by subtracting the expected phenotype from the measured phenotype of those combined knockdowns.

Ribosome Profiling:

K562-dCas9 cell line stably expressing $GFP_{Non\text{-}stop}$ was infected with lenti vectors containing a guide RNA against one gene or non-targeting guide RNA (Table 2). Cells were selected using FACS to recover cells containing sgRNA (labeled with RFP). Cyclohexamide was added (100mg/ul) to the cells and they were incubated for 5 minutes at 37 °C and then pelleted at 400g for 5 minutes. Media was removed and cell pellets were immediately snap frozen. Libraries were prepared according to McGlincy and Ingolia, Methods, 2017.

RNA sequencing:

RNA was extracted using Zymo mini RNA prep (R2053) and libraries were cloned using Illumnia true stranded total RNA seq kit (20020596).

Ribosome profiling and RNAseq analysis:

Ribosome profiling and RNA-seq libraries were sequenced on an Illumina HiSeq 4000 as single end 50-bp reads (UCSF Center for Advanced Technology) at a depth of 22-64 million reads per library. Ribosome profiling reads were processed by removing 3' linker sequences using *FASTX clipper* and de-multiplexed using *FASTX barcode splitter* (http://hannonlab.cshl.edu/fastx_toolkit/). Sample barcodes and unique molecular identifiers were trimmed using a custom Python script. Next, abundant reads were filtered by aligning to a custom library of human rRNAs, tRNAs, and primer sequences using *bowtie* v1.2 (http://bowtie-bio.sourceforge.net/). All remaining reads were aligned to a modified human genome (GRCh38.92 with addition of our CRISPRi construct and the $GFP_{Non\text{-}stop}$) using *tophat* v2.1.1 (https://ccb.jhu.edu/software/tophat/; relevant parameters: --read-mismatches=1 and --no-novel-juncs). After alignment, uniquely mapping reads were extracted, and *plastid* (https://plastid.readthedocs.io) was used to obtain gene-level read counts and normalized counts, reads per kilobase per million mapped reads (RPKM). To assign reads to genomic positions, a p-site offset of 12 nucleotides was determined by calculating the average distance of the start codon peak from the start codon (*plastid psite*). To assess the quality of the ribosome profiling data, we confirmed that the aligned reads displayed the expected three nucleotide periodicity (phasing) and that the majority of ribosome footprints had a length of 28-31 bp for all libraries. We also confirmed that the CRISPRi system produced robust target knockdown on the day of the experiment at the level of ribosome footprints (knockdown of target genes >88% for all genes). The RNA-seq libraries were processed using the same pipeline as the ribosome profiling libraries, with the exception that *tophat* was run with –read-mismatches=2 due to the longer RNAseq read length. All downstream analyses were performed using the Python libraries pandas, plastid, matplotlib, seaborn, numpy, and scipy.

For Figure S1A, footprints from K562 CRISPRi cells bearing non-targeting sgRNA are plotted as reads per million. Here, we plotted all aligned reads rather than uniquely aligned reads to visualize read density in the $GFP_{Non\text{-}stop}$ MALAT1 triple helix, which is identical to the endogenous MALAT1 lncRNA. To confirm that the reads aligning to the MALAT1 triple helix in the $GFP_{Non\text{-}stop}$ are not derived from the endogenous MALAT1 lncRNA, we performed ribosome profiling in a K562 CRISPRi cell line without the $GFP_{Non\text{-}stop}$ reporter. Here, we found that the endogenous MALAT1 background rate was <10% of the $GFP_{Non\text{-}stop}$ footprints in the MALAT1 triple helix, suggesting that the $GFP_{Non\text{-}stop}$ triple helix is indeed translated. For Figure S5A, translational efficiencies were calculated by normalizing the $GFP_{Non\text{-}stop}$ GFP ribosome footprint rpm to the $GFP_{Non\text{-}stop}$ RNAseq rpm.

RT-qPCR:

Total RNA was isolated from frozen cell samples using TRIzol reagent (Thermo Fisher Scientific) and Phase Lock Gel tubes (VWR), treated with Turbo DNase (Thermo Fisher Scientific), or using Direct-zol RNA MiniPrep kit. Reverse-transcription was carried using M-MLV (Thermo Fisher Scientific) or SSIII

Reverse-transcriptase (Thermo Fisher Scientific) with random hexamer primers (Thermo Fisher Scientific, SO124) in the presence of RNaseIN Recombinant Ribonuclease Inhibitor (Thermo Fisher Scientific). Quantitative PCR (qPCR) was performed with Kappa Sybr Fast qPCR 2x Mix (Roche), according to the manufacturer's instructions on a LightCycler 480 Instrument (Roche). Experiments were performed in technical triplicates. RT-qPCR primers used are listed in (Table 3).

Immunoprecipitation and western blot:

Cells were lysed in buffer containing 10mM Tris 7.5, 150 mM NaCl, 0.5% NP-40, 5 mM $Mg_2Cl$, and 1x Halt™ Protease Inhibitor Cocktail (Thermo Fisher Scientific). The lysates were cleared by centrifugation at 20,000 x g for 10 min and bound to GFP-Trap Magnetic Beads (Chrometech) for 2h at 4 °C. Beads were washed 3 times with IP Buffer (10mM Tris 7.5, 150 mM NaCl, 0.1% NP-40, 5 mM $Mg_2Cl$). Bound material was eluted by boiling in Laemmli Buffer for 10 min at 90°C.

Proteins were separated on Bolt® 4-12% Bis-tris gels (Thermo Fisher Scientific), transferred to PVDF membrane using the Mini Trans-Blot Cell (Bio-Rad) according to the manufacturer's instructions, blocked with 5% milk in TBS, and subsequently probed. Rabbit anti ZNF598 antibody (Bethyl Laboratories, A305-108A-T), rabbit anti GIGYF2 antibody (Bethyl Laboratories, A303-731A-M), and rabbit anti 4EHP antibody (Cell Signaling, 6916S) primary antibodies were used. LI-COR IRDye800/IRDye700 anti-rabbit (Odyssey) secondary antibody was used at 1:10,000 dilution. All blots were visualized using the LI-COR (Odyssey) system.

ZNF598 Mass Spectrometry:

*Cell lines*

3X-FLAG-tagged ZNF598 overexpressing (ZNF598-OE) Flp-In T-Rex 293 cells (kind gift from Dr. Nahum Sonenberg and Dr. Thomas Tuschl, Garzia *et al.*, Nature Communications, 2017) were grown in high-glucose Dulbecco's Modified Eagle Medium (Thermo Fisher, 11995040, containing 4 mM L-Glutamine and 1 mM sodium pyruvate) supplemented with 10% (v/v) fetal bovine serum. The affinity purification mass spectrometry experiment was performed in biological triplicate. Two million low-passage cells were seeded per 10 cm dish. Expression of 3X-FLAG-tagged ZNF598 was induced (24 hours post seeding) for 24 hours by the addition of doxycycline at 1 μg/ml at final concentration.

*FLAG-immunoprecipitation of 3X-FLAG-ZNF598 from HEK293 cells*

Cells were washed with warm PBS (containing 360 μM emetine), and harvested in ice-cold lysis buffer (50 mM TRIS pH 8.0, 150 mM KCl, 15 mM $MgCl_2.6H_2O$, 5% glycerol) supplemented with 1% Triton X-100, 360 μM emetine, 20 mM N-ethylmaleimide, 80 units TURBO DNase (Thermo Fisher, AM2238) and protease inhibitors (cOmplete Protease Inhibitor Cocktail, Roche; Protease Inhibitor Cocktail, Millipore Sigma; and PMSF, Millipore Sigma). Cell lysates were clarified in a tabletop micro-centrifuge (15 minutes, 4°C). Equal protein amounts from the clarified supernatant were used for each immunoprecipitation. The clarified supernatant was incubated with ANTI-FLAG-Agarose M2 affinity gel (Millipore Sigma #A2220, 15 µl of packed affinity resin) for 1 hour, washed four times with 400 µl wash 1 buffer (lysis buffer supplemented with 0.1% Triton X-100 and 360 µM emetine), then washed four times with wash 2 buffer (50 mM TRIS pH 8.0, 150 mM KCl, 15 mM $MgCl_2.6H_2O$ supplemented with 360 µM emetine), and eluted in 80 µl elution buffer (50 mM TRIS pH 8.0, 150 mM KCl, 15 mM $MgCl_2.6H_2O$ supplemented with 400 µg/ml 3X FLAG-peptide (Millipore Sigma, #F4799)). Eluates were flash frozen in liquid nitrogen and stored at -80°C prior to protein digestion.

*Protein Digestion*

Protein extracts (4 µg) were diluted up to 300 µl in 10 mM triethyl ammonium bicarbonate (TEAB) and were reduced with 15 µl of 7.5 mg/ml DL-dithiothreitol (DTT) (60°C, 1 hour). After cooling to room temperature, samples were alkylated with 15 µl of 18.5 mg/ml iodoacetamide for 15 minutes at room temperature in the dark. Reduced and alkylated proteins were buffer-exchanged on a 30 kDa molecular weight spin cartridge (Amicon Ultra 0.5 ml, Millipore Sigma) and washed four times with 400 µl 10 mM TEAB. Proteins were digested overnight at 37°C on the filter with 300 µl Trypsin (20 µg in 3 ml 10 mM TEAB, Promega Sequencing Grade Modified Trypsin). Additional Trypsin (100 µl of 10 mg/ml) was added the next morning (37°C, 1 hour). Peptides were removed from the top of the filter and the filter was washed twice with 300 2% acetonitrile, 0.1% formic acid. All washes were combined and dried.

*Liquid Chromatography and Mass Spectrometry*

Liquid Chromatography and Mass Spectrometry was performed at the Mass Spectrometry and Proteomics Core, JHMI. Peptides were analyzed by liquid chromatography interfaced with tandem mass spectrometry (LC/MS/MS) using a Waters nanoACQUITY UPLC system (www.waters.com) interfaced with an Orbitrap Fusion™ Lumos™ Tribrid™ Mass Spectrometer (www.thermofisher.com). Fractions were resuspended in 20 µl loading buffer (2% acetonitrile in 0.1% formic acid) and analyzed by reverse phase liquid chromatography coupled to tandem mass spectrometry. Peptides (50%, approx. 1 µg) were loaded onto a C18 trap (S-10 µM, 120 Å, 75 µm x 2 cm, YMC, Japan) and subsequently separated on an in-house packed PicoFrit column (75 µm x 200 mm, 15u, +/-1 µm tip, New Objective) with C18 phase (ReproSil-Pur C18-AQ, 3 µm, 120 Å, www.dr-maisch.com) using 2-90% acetonitrile gradient at 300 nl/min over 120 min. Eluting peptides were sprayed at 2.0 kV directly into the Lumos.

Survey scans (full MS) were acquired from 350-1800 m/z with data-dependent monitoring with a 3 sec cycle time. Each precursor individually isolated in a 1.6 Da window and fragmented using HCD activation collision energy 30 and 15 sec dynamic exclusion, first mass being 120 m/z. Precursor and fragment ions were analyzed at resolutions 120,000 and 30,000, respectively, with automatic gain control (AGC) target values at 4e5 with 50 ms maximum injection time (IT) and 1e5 with 100 ms maximum IT, respectively.

*Processing of Mass Spectrometry Data*

Raw data was processed and analyzed using the MaxQuant software suite (Cox and Mann, Nature Biotechnology, 2008). Default settings were used except that 'Match between runs' was turned on to transfer peptide identification from an LC-MS run, in which the peptide has been identified by MS/MS, to another LC-MS run, in which no MS/MS data for the same peptide was acquired or no peptide was assigned (Tyanova *et al.*, Nature Methods, 2016). Search parameters were as follows:

a maximum of two missed cleavages were allowed, cysteine carbamidomethyl was included as a fixed modification, and variable modifications included oxidation of methionine, protein N-terminal acetylation, deamidation of glutamine and asparagine, phosphorylation of serine, threonine and tyrosine, and Gly-Gly ubiquitin remnant on lysine. Database search was performed with Andromeda (Cox and Mann, Nature Biotechnology, 2008; Cox et al., Journal of Proteomic Research, 2011) against Uniprot human database (UP000005640_9606.fasta; downloaded on 09/10/2018) with common serum contaminants and enzyme sequences. False discovery rate (FDR) was set to 1% at both the peptide spectrum match (PSM) and protein level. A minimum peptide count required for protein quantification was set to two. Protein groups were further analyzed using the Perseus software suite (Tyanova *et al.*, Nature Methods, 2016). Common contaminants, reverse proteins and proteins only identified by site were filtered out. LFQ values were transformed to $log_2$ space and intensity distributions were checked to ensure that data was normally distributed.

**Supplemental Figures:**

Hickey *et al.*, Fig. S1

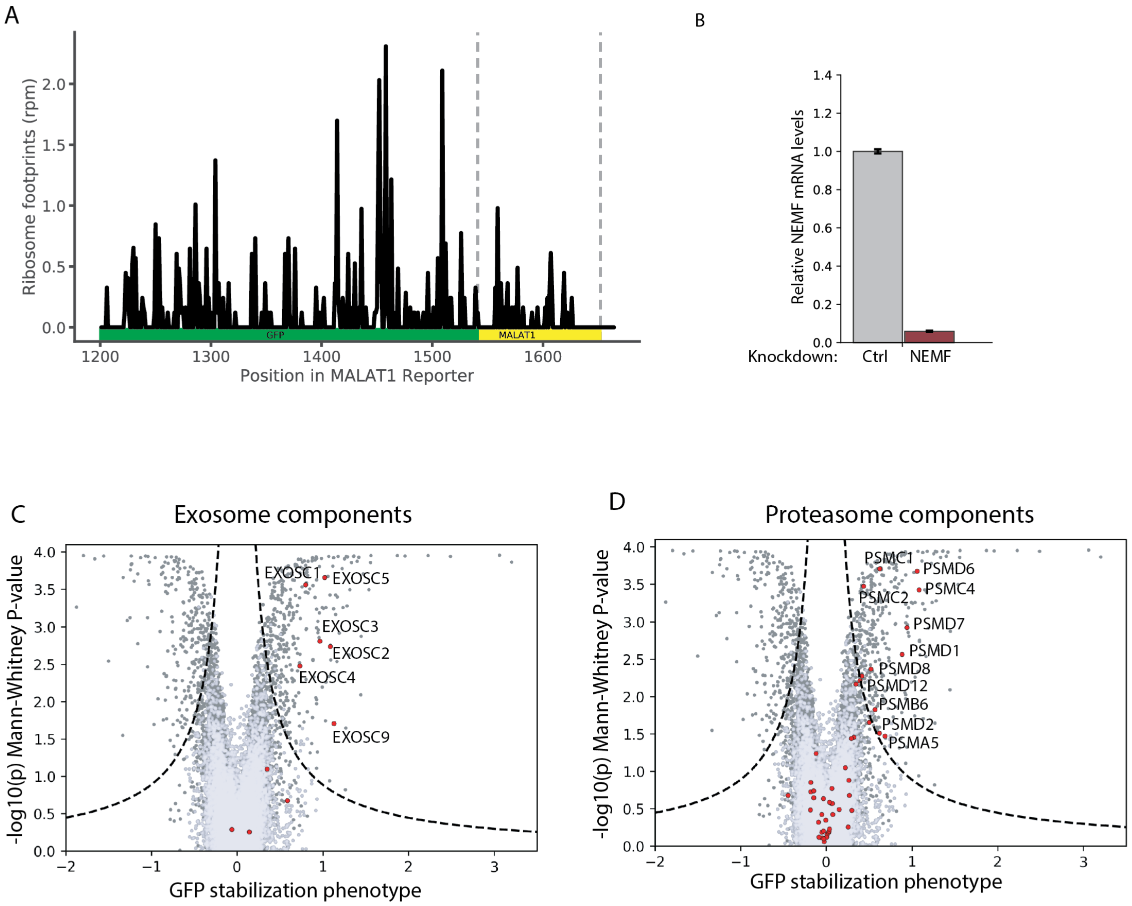


**Figure S1. GFP$_{\text{Non-stop}}$ is a non-stop stalling reporter. (A)** Ribosome occupancy (RPM) along the GFP$_{\text{Non-stop}}$ reporter. (**B**) Relative *NEMF* mRNA levels measured by qPCR (median ± SD, N=3). (**C, D**) FACS screen volcano plot with Exosome or proteasome components highlighted in red.

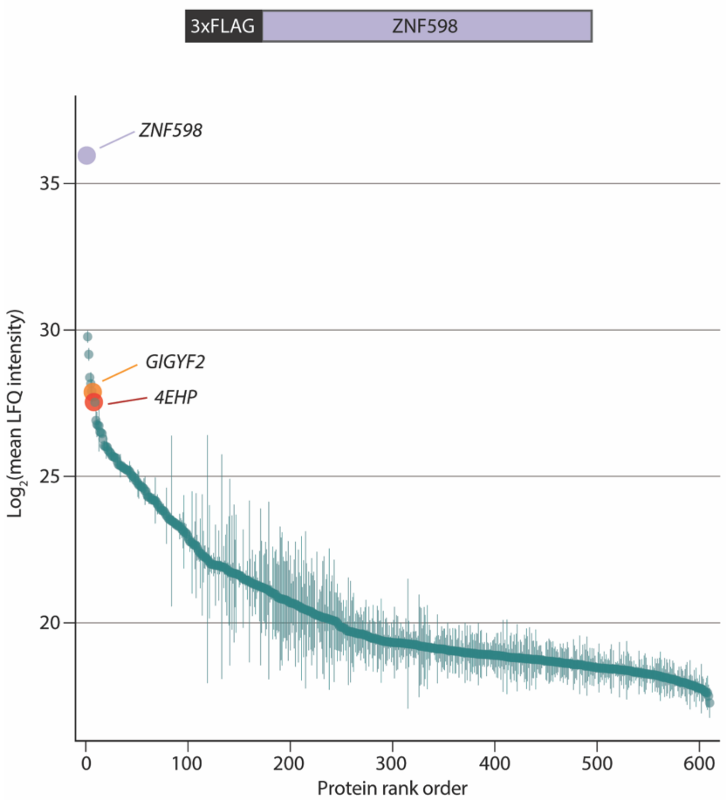


**Figure S2. GIGYF2 and 4EHP are top interactors of ZNF598.** Ranked proteins identified via mass-spectrometry after 3FLAG-ZNF598 immunoprecipitation (mean ± SD, N=3)

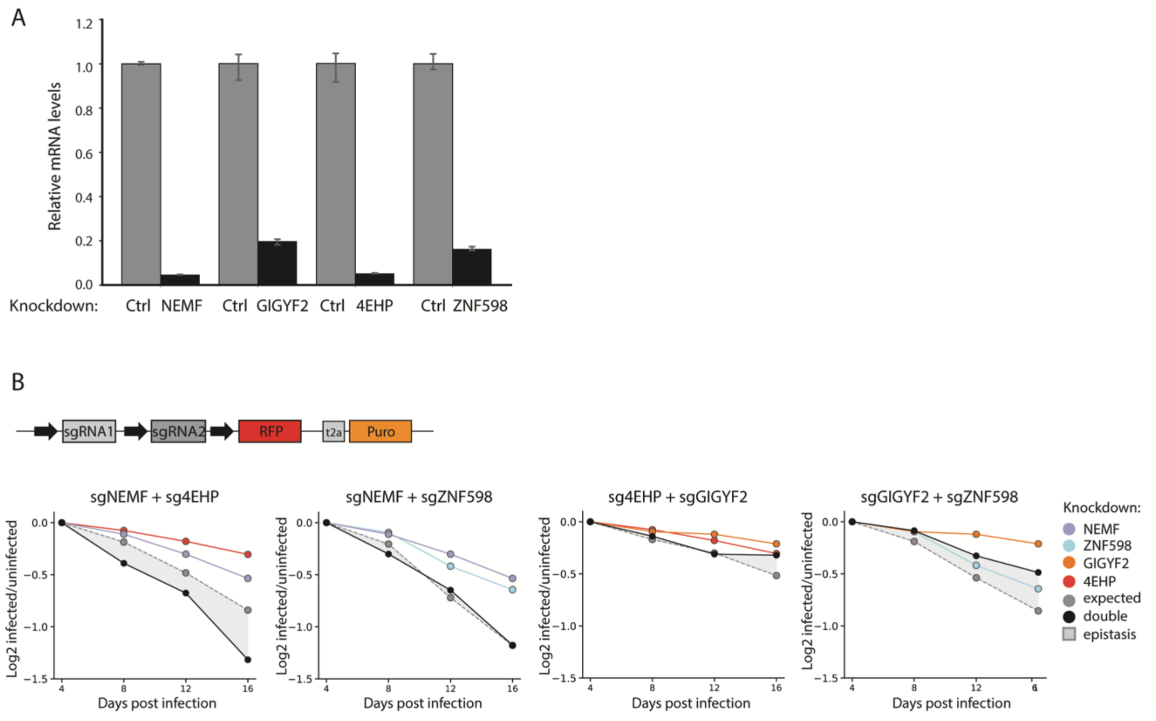


**Figure S3. GIGYF2 and 4EHP have a synergistic growth interaction with NEMF.** (**A**) Relative mRNA levels for *NEMF, GIGYF2, 4EHP,* and *ZNF598* upon knockdown measured by qPCR (mean ± SD, N=3). (**B**) Competition assays between control cells and cells harboring a single or double knockdown of *NEMF*, ZNF598, *GIGYF2,* and *4EHP*.

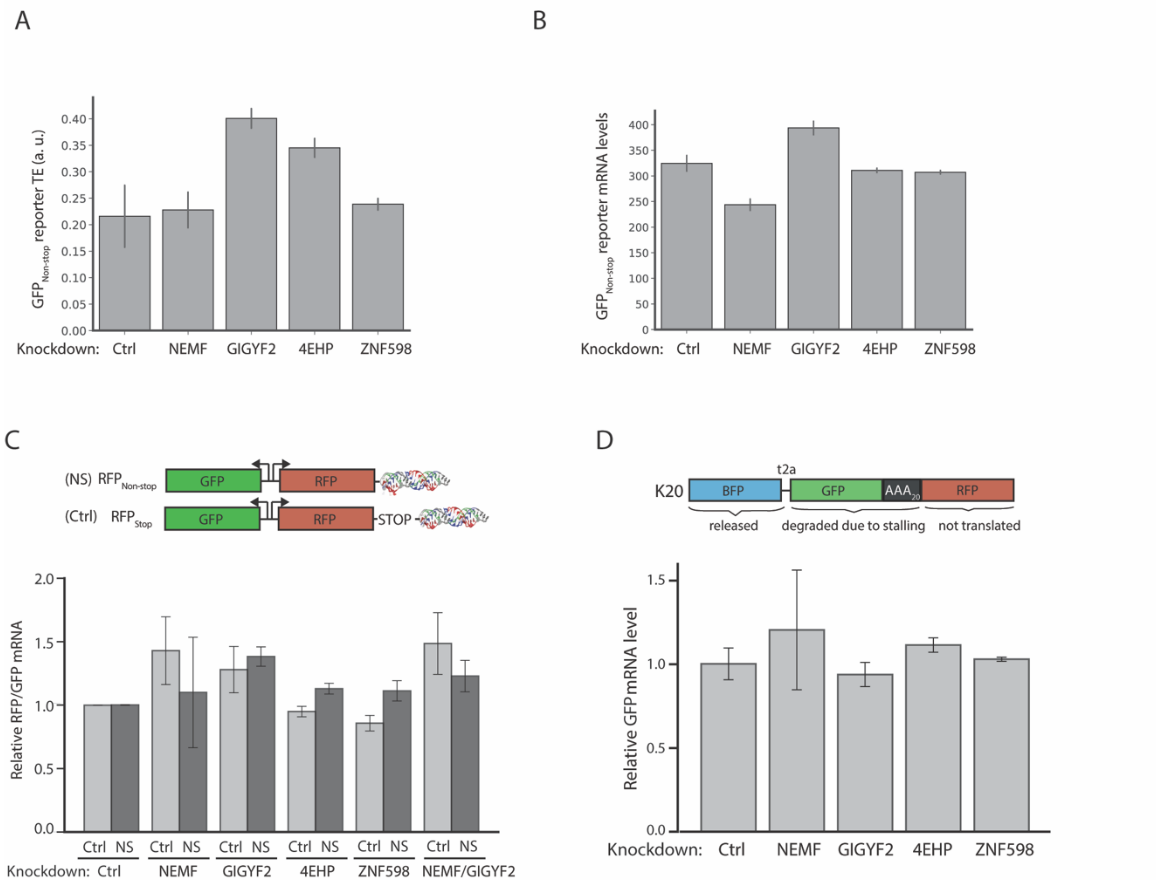


**Figure S4. GIGYF2 and 4EHP decrease the ribosome load on faulty mRNAs.** (**A**) Translation efficiency (ribosome profiling footprints normalized to mRNA levels) of $GFP_{Non\text{-}stop}$ reporter in knockdown cell lines. (**B**) mRNA levels of $GFP_{Non\text{-}stop}$ reporter in knockdown cell lines. (**C**) RFP/GFP mRNA levels for $RFP_{Non\text{-}stop}$ and $RFP_{Stop}$ reporters in knockdown cell lines. (**D**) Relative mRNA levels for K20 stalling reporter in knockdown cell lines.

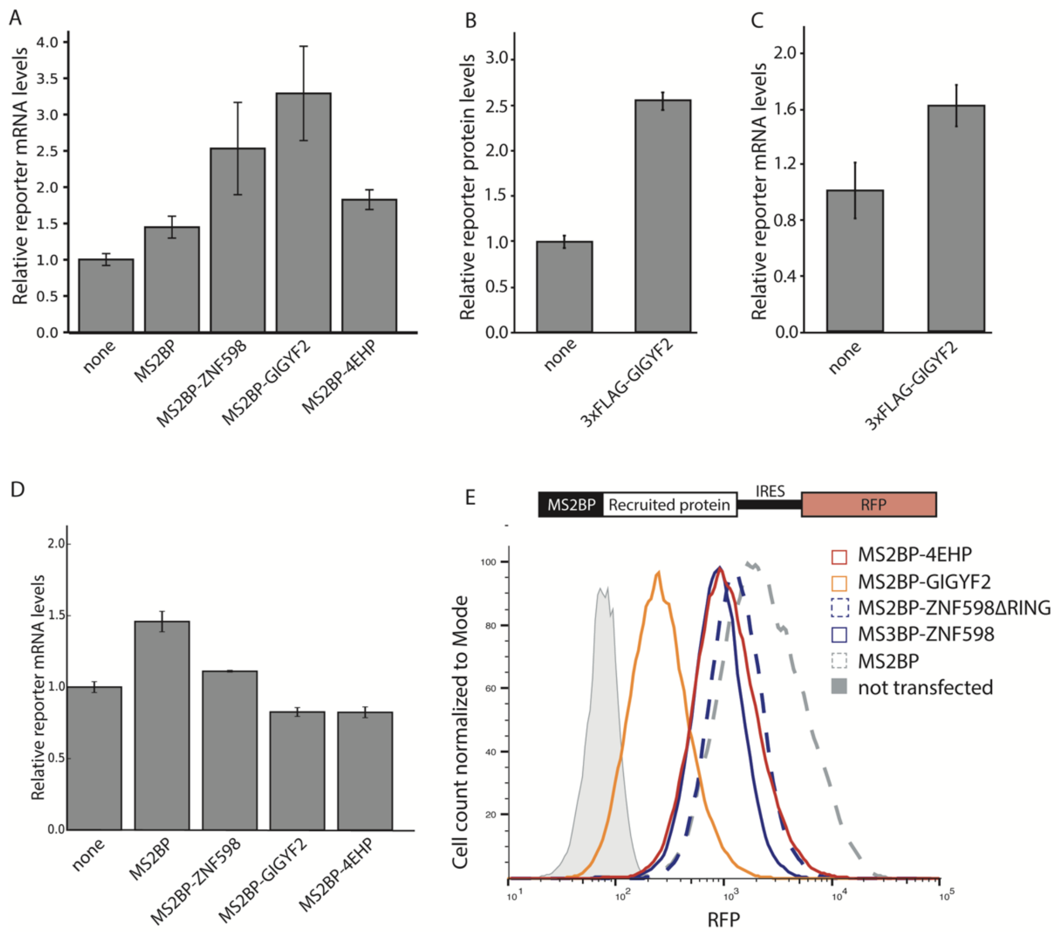


**Figure S5. GIGYF2, 4EHP, and ZNF598 inhibit translation when directly recruited to an mRNA.** (**A, D**) Relative mRNA levels of $GFP_{MS2\text{-}Stop}$ reporter upon transient or stable expression of MS2BP-fusion proteins (mean ± SD, N=3). (**B, C**) Protein and mRNA levels of $GFP_{MS2\text{-}Stop}$ measured by flow cytometry or qPCR upon transient expression of 3xFLAG-GIGYF2, that is not directly recruited to the reporter (mean ± SD, N=3). (**E**) MS2BP-fusion proteins expression levels measured by RFP fluorescence.

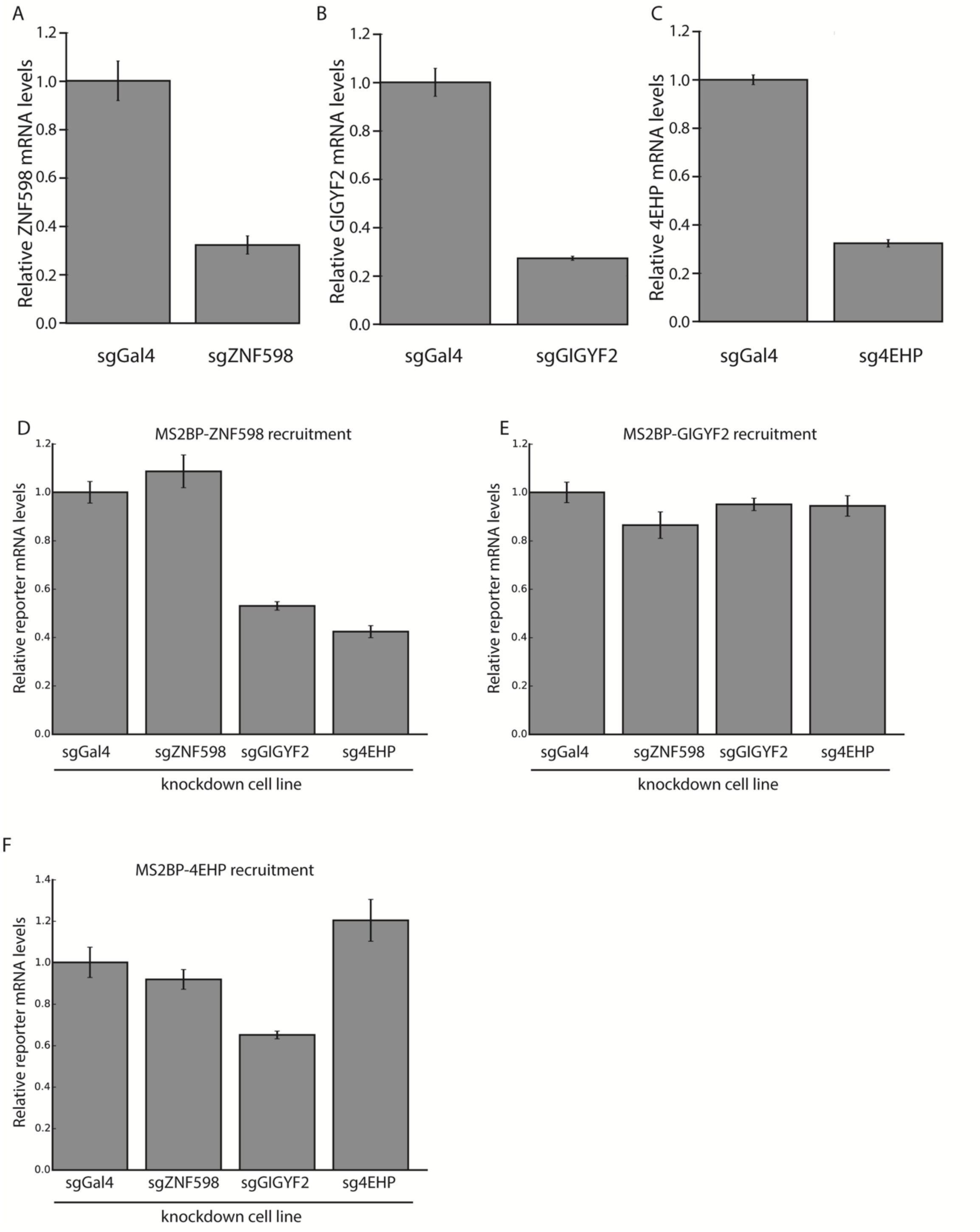


**Figure S6. ZNF598 mediated translation inhibition requires GIGYF2 and 4EHP.** (**A, B, C**) Relative mRNA levels upon control (Gal4) or targeting (ZNF598, GIGYF2, 4EHP) knockdown in cells stably expressing $GFP_{MS2\text{-}Stop}$ reporter (mean ± SD, N=3). (**D, E, F**) Relative $GFP_{MS2\text{-}Stop}$ reporter mRNA levels in knockdown cell lines upon MS2-fusion protein recruitment (mean ± SD, N=3).

Hickey *et al.*, Fig. S7

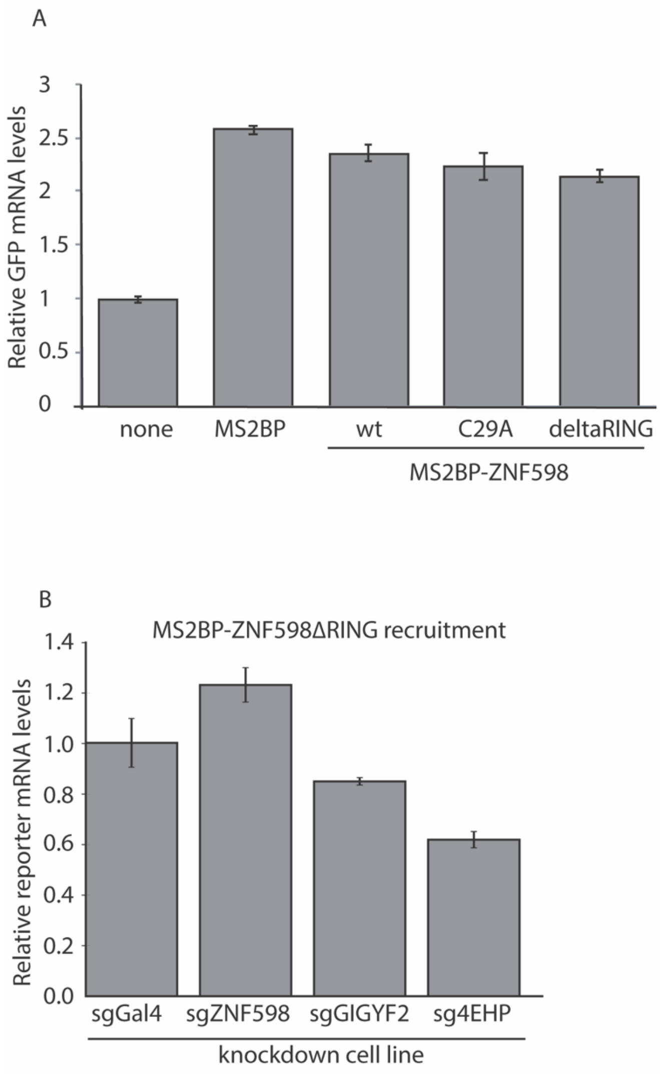


**Figure S7. ZNF598 mediated translation inhibition is ubiquitination independent.** (**A**) Relative mRNA levels of $GFP_{MS2\text{-}Stop}$ reporter upon MS2BP-ZNF598 (wt, C29A, ΔRING) recruitment (mean ± SD, N=3). (**B**) Relative mRNA levels of $GFP_{MS2\text{-}Stop}$ reporter in knockdown cell lines upon MS2BP-ZNF598 (wt and ΔRING) fusion protein recruitment (mean ± SD, N=3).

**Supplemental Tables:**

**Table 1: Plasmids used in this study**

| Name | ID | Backbone | Expression | Source |
|---|---|---|---|---|
| GFP Non-stop | pKK100 | pcDNA3 | pCMV-BFP-t2a-GFP-Malat1 | this study |
| GFP Non-stop | pKK122 | piggybac | pEIF2a-BFP-T2A-GFP-Malat1 | this study |
| GFP Stop | pKK357 | pcDNA3 | pCMV-BFP-t2a-GFP-STOP-Malat1 | this study |
| GFP PolyA | pKK358 | pcDNA3 | pCMV-BFP-t2a-GFP-STOP-polyA | this study |
| K20 | pKH21 | piggybac | pEIF2a-BFP-T2A-GFP-K20-RFP | this study |
| RFP Non-stop | pKH63 | piggybac | GFP-pA; RFP-MALAT1 | this study |
| RFP Stop | pKH64 | piggybac | GFP-pA; RFP-STOP-MALAT1 | this study |
| yeast control reporter | | | RFP-pA; GFP-T2A-FLAG-His3 | D'Orazio et al., eLife 2019 |
| yeast NGD reporter | | | RFP-pA; GFP-T2A-FLAG-His3(CGA)12 | D'Orazio et al., eLife 2019 |
| GFP MS2-Stop | pKK306 | pcDNA | pCMV-3xMS2_BFP-t2a-GFP-stop-stop-Malat1 | this study |
| GFP MS2-Stop | pKK307 | piggybac | pEIF2a-3xMS2_BFP-IRES-GFP-stop-stop-Malat1 | this study |
| MS2BP | pKK308 | pcDNA | MS2BP-IRES-mCherry | this study |
| MS2BP-GIGYF2 | pKK309 | pcDNA | MS2BP-GIGYF2-IRES-mCherry | this study |
| MS2BP-ZNF598 | pKK375 | pcDNA3 | MS2BP-ZNF598-IRES-mCherry | this study |
| MS2BP-4EHP | pKK318 | pcDNA3 | MS2BP-4EHP-IRES-mCherry | this study |
| MS2BP-ZNF598_C29A | pKK391 | pcDNA3 | MS2BP-ZNF598_C29A-IRES-mCherry | this study |
| MS2BP-ZNF598_deltaRING | pKK392 | pcDNA3 | MS2BP-ZNF598_deltaRING-IRES-mCherry | this study |
| MS2BP | pKK478 | piggybac | MS2BP-IRES-Mcherry | this study |
| MS2BP-ZNF598_deltaRING | pKK479 | piggybac | MS2BP-ZNF598deltaRING-IRES-mCherry | this study |
| MS2BP-GIGYF2 | pKK480 | piggybac | MS2BP-GIGYF2-IRES-mCherry | this study |
| MS2BP-4EHP | pKK481 | piggybac | MS2BP-4EHP-IRES-mCherry | this study |
| MS2BP-ZNF598 | pKK482 | piggybac | MS2BP-ZNF598WT-IRES-mCherry | this study |

**Table 2: Protospacers used in this study**

| Target | sequence |
|---|---|
| non-targeting 001 | GAACGACTAGTTAGGCGTGTAG |
| NEMF | GAGTACGGCGCGGAGGTCAA |
| GIGYF2 | GGCAGGGGAGCGACACGGAA |
| 4EHP | GCCTGAGGCAGTGGCGACAG |
| ZNF598 | GGCCGGATCCCGGACCATGG |
| non-targeting 002 | GTGTGCAACCTCCGCCGTTGG |

**Table 3: RT-qPCR primers used in this study**

| Target | forward primer | reverse primer |
|---|---|---|
| ZNF598 | aaaggtgtacgcattgtacagg | ctccaggtccccgaagag |
| 4EHP | gacaggccacagtgacttcc | gcagccgaataatccacttg |
| GIGYF2 | tctgtgggtcaggaatttgg | gacatctgaccacaaccaaaga |
| Actin | gctacgagctgcctgacg | ggctggaagagtgcctca |
| GFP | tgaccacatggtccttcttg | atagttcatccatgccatgtgta |
| GAPDH | acgggaagcttgtcatcaat | catcgccccacttgatttt |
| RFP | cccgtaatgcagaagaagacc | gcttgatctcgcccttca |
| copGFP | gtggatggcgctcttgaa | cgacggcggctactacag |
| NEMF | ctatatgaaaacaacatccaacttcag | cgtcagtatgtcttcaactggttt |
